# Biophysical demonstration of co-packaging of glutamate and GABA in individual synaptic vesicles in the central nervous system

**DOI:** 10.1101/2021.03.23.436594

**Authors:** SeulAh Kim, Michael L. Wallace, Mahmoud El-Rifai, Alexa R. Knudsen, Bernardo L. Sabatini

## Abstract

Many mammalian neurons release multiple neurotransmitters to activate diverse classes of ionotropic receptors on their postsynaptic targets. Entopeduncular nucleus somatostatin (EP *Sst+*) neurons that project to the lateral habenula (LHb) release both glutamate and GABA, but it is unclear if these are packaged into the same or segregated pools of synaptic vesicles. Here we describe a novel method combining electrophysiology, spatially-patterned optogenetics, and computational modeling designed to analyze the mechanism of glutamate/GABA corelease. We find that the properties of PSCs elicited in LHb neurons by optogenetic activation of EP *Sst+* terminals are only consistent with co-packaging of glutamate and GABA into individual vesicles. Furthermore, serotonin, which acts presynaptically to weaken EP *Sst+* to LHb synapses, does so by altering the release probability of vesicles containing both transmitters. Our approach is broadly applicable to the study of multi-transmitter neurons throughout the brain and our results constrain mechanisms of neuromodulation in LHb.

## Introduction

Many neurons in the mammalian brain can produce, store and release multiple neurotransmitters (Tritsch et al., 2016). Co-release refers to chemical release of two or more neurotransmitters, emphasizing a property of the pre-synaptic terminal. In contrast, the term co-transmission highlights the functional aspect of neurotransmission, hence implying the presence of post-synaptic receptors that detect each of the released transmitters. Despite the prevalence of multi-transmitter neurons throughout the brain, our understanding of how, when, and where multiple neurotransmitters are released and what purpose such co-release serves remains incomplete.

The mechanisms and post-synaptic consequences of neurotransmitter co-release from multi-transmitter neurons varies. For example, in some cases multiple small-molecule (i.e. non-peptide transmitters) neurotransmitters are thought to be packaged into the same vesicle (Jonas et al., 1998; Shabel et al., 2014; Tritsch et al., 2012) whereas in other cases a single cell makes multiple classes of pre-synaptic boutons, each of which releases a different transmitter (Granger et al., 2020; Lee et al., 2010; Zhang et al., 2015). Furthermore, even if two transmitters are released in the same vesicle from a single synaptic bouton, the opposing post-synaptic target may not have receptors for both transmitters, preventing co-transmission of the signal to the post-synaptic cell. Conversely, two transmitters may be released from different presynaptic terminals, but, if these are made onto the same post-synaptic cell, co-transmission will occur. For these reasons, it is technically challenging to functionally analyze the mechanisms of neurotransmitter co-release and reveal their importance to neural circuits. In particular, the mechanisms of co-release and co-transmission at synapses formed by multi-transmitter neurons is difficult to determine from the average synaptic responses, necessitating experiments examining single release events from single synapses.

Co-transmitting neurons are found in the entopeduncular nucleus (EP), a basal ganglia output nucleus comprised of multiple neural populations differentiable by their transcriptome, the types of neurotransmitters they release, and their projection targets. Somatostatin positive (*Sst+*) EP neurons project solely to the lateral habenula (LHb) and express the molecular machinery necessary to release glutamate and GABA (Wallace et al., 2017). Indeed, stimulation of EP *Sst+* axons causes release of glutamate and GABA and results in compound synaptic currents in postsynaptic LHb neurons mediated by opening of ionotropic glutamate and GABA receptors (Root et al., 2018; Wallace et al., 2017).

Although individual axons of *Sst+* EP neurons are thought to release both glutamate and GABA, their mechanism of co-transmission remains inconclusive. One proposed mechanism is co-packaging of glutamate and GABA in the same vesicles. This model is supported by the detection of biphasic miniature spontaneous synaptic responses in LHb neurons, suggesting that they are generated by glutamate and GABA co-released from individual vesicles (Shabel et al., 2014). A second model is segregation of glutamate and GABA into different pools of synaptic vesicles that are independently released from the same terminal. This model is supported by ultrastructural evidence showing that the glutamate and GABA vesicular transporters, Vglut2 and Vgat, respectively, are found in separate pools of vesicles within the same axon terminals in LHb (Root et al., 2018). Moreover, synaptic vesicles isolated from LHb are immunoreactive against either Vgat or Vglut2 (Root et al., 2018).

Whether glutamate and GABA release from EP *Sst+* neurons in the LHb occurs via co-packaging in individual vesicles or by co-transmission from separate pools has important functional consequences. LHb regulates major monoaminergic centers in the brain (Hu et al., 2020; Matsumoto and Hikosaka, 2009, 2007). EP heavily innervates the lateral portion of LHb and is implicated in aversion, encoding of reward prediction error and action-outcome evaluation (Hong and Hikosaka, 2008; Li et al., 2019; Shabel et al., 2012; Stephenson-Jones et al., 2016). Furthermore, synaptic plasticity that shifts the relative proportion of glutamatergic vs. GABAergic co-transmission from EP to LHb alters the excitability (Li et al., 2011) and bursting states of LHb neurons (Yang et al., 2018). This change is thought to drive animals towards maladaptive behavior states, such as depression, chronic-stress induced passive coping, and addiction (Cerniauskas et al., 2019; Li et al., 2011; Maroteaux and Mameli, 2012; Meye et al., 2016; Shabel et al., 2014; Trusel et al., 2019). Therefore, the mechanism by which glutamate/GABA co-transmission occurs, and how it may be modulated by plasticity, likely has important functional implications for stress, anxiety and depression.

Here we combine molecular, computational, pharmacological and electrophysiological analyses to distinguish the two models of glutamate and GABA co-release at synapses between EP *Sst+* and LHb neurons. Immunohistochemical analysis of the distributions of synaptic proteins reveals that the proteins necessary for glutamate and GABA release are colocalized within individual EP *Sst+* terminals. We characterize differential statistical features expected by the two distinct release modes and compare them to experimental results collected using an advanced optogenetic activation approach that targets individual EP *Sst+* boutons. We discover that glutamate and GABA are co-packaged in the same vesicles in EP *Sst+* terminals. In addition, serotonin co-modulates release of both glutamate and GABA while maintaining the correlation between glutamatergic and GABAergic unitary responses, further supporting that the two transmitters are released from the same vesicle. Our methods are generally applicable to the study of the mechanism of co-release of neurotransmitters from multi-transmitter neurons. Our findings have important implications for plasticity mechanisms underlying shifted balance of glutamatergic and GABAergic transmissions between EP and LHb in maladaptive states.

## Results

### Functional and molecular evidence of co-release of glutamate and GABA from EP *Sst+* axons in LHb

Somatostatin-expressing neurons (*Sst+*) that reside in the anterior region of the EP release both glutamate and GABA (Shabel et al., 2014; Wallace et al., 2017). To gain optogenetic control, we replicated a previous approach that transduces *Sst+* neurons’ cell bodies in the EP and labels their axons in the LHb (Wallace et al., 2017). We bilaterally injected adeno-associated virus (AAV) that expresses the channelrhodopsin variant oChIEF in a Cre-dependent manner (AAV-DIO-oChIEF) into the EP of *Sst-IRES-Cre* (*Sst-Cre*) mice (Figure 1A) (Lin et al., 2009; Taniguchi et al., 2011). Consistent with a previous study demonstrating monosynaptic release of glutamate and GABA (Wallace et al., 2017), optogenetic activation triggered a biphasic post-synaptic current (PSC) in a LHb neuron under whole-cell voltage-clamp recording (holding voltage, V_h_ = −35 mV) in the presence of the NMDA receptor antagonist CPP (Figure 1B). This current profile results from the faster opening and closing kinetics of AMPA receptors (AMPARs) compared to GABA_A_ receptors (GABA_A_Rs). Both GABAergic and glutamatergic currents persist in the presence of TTX/4AP, consistent with direct release of both transmitters from the optogenetically stimulated axons (Wallace et al., 2017). Thus, this genetic strategy grants access to and permits manipulation of the glutamate/GABA co-releasing EP-to-LHb projections.

**Figure 1.**
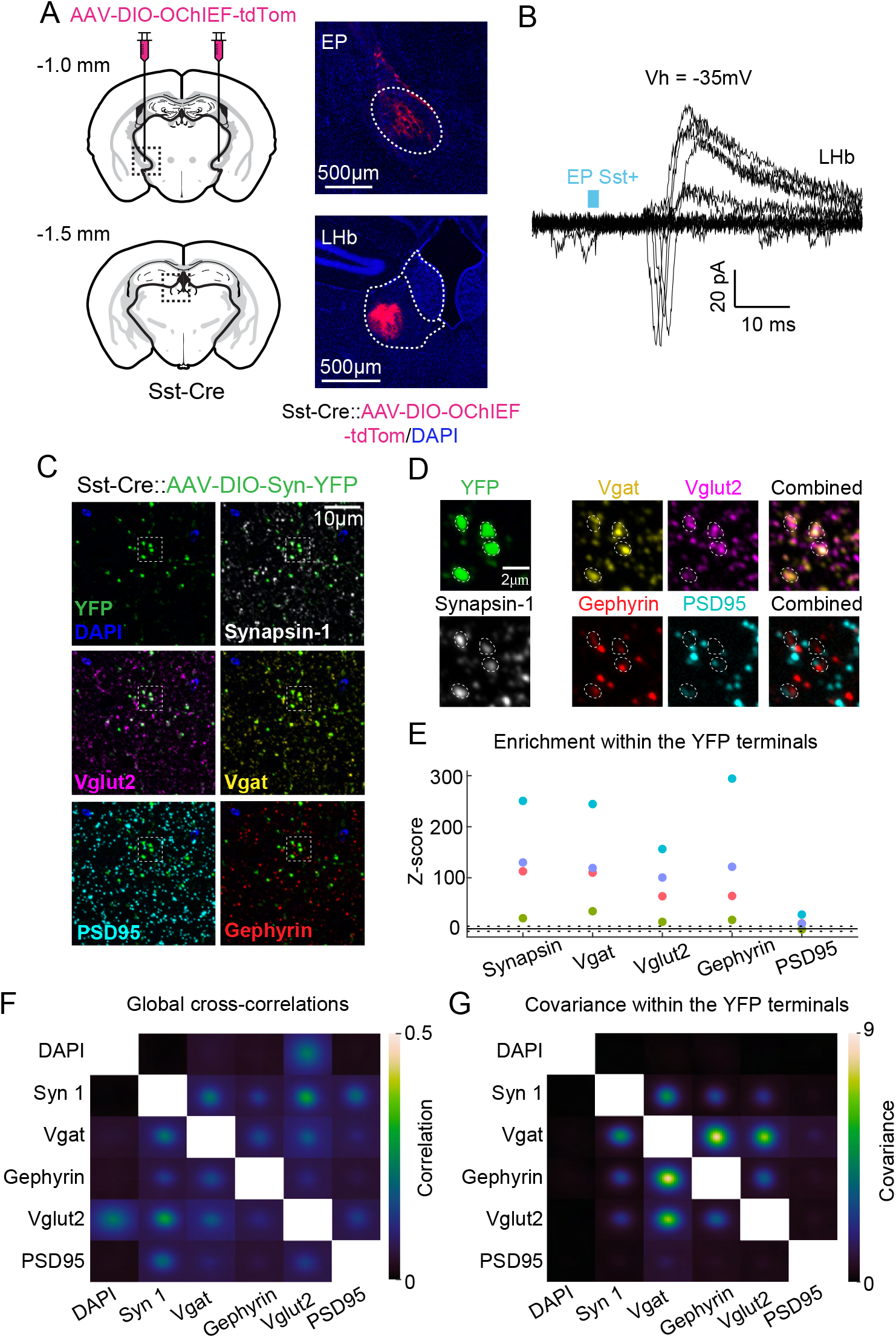
Electrophysiological and molecular evidence for glutamate and GABA co-release from EP Sst+ axons in LHb. A) *left*, Schematic of the experimental design. *Sst-Cre* mice were bilaterally injected with Cre-dependent AAV-DIO-OChIEF-tdTom into the EP (*top*) resulting in axonal labeling of projections to the LHb (*bottom*). *right*, Histological analysis showing expression of tdTom in cell bodies at the injection site (*top*) and in axons of EP *Sst+* projection neurons in the LHb (*bottom*). Scale bars=500 µm B) Example post-synaptic currents (PSCs) recorded from a LHb neuron (V_h_=-35 mV) following optogenetic activation of EP *Sst+* axons using wide-field minimal photo stimulation in an acute brain slice. With repetitive stimulation at minimal power, some trials result in failure of release whereas other trials lead to successful release events that evoke both inward and outward currents as seen in the biphasic PSCs. The blue box shows the timing and duration of the laser pulse. C) Serial sections of brain tissue containing EP *Sst+* axon terminals labeled with Synaptophysin-YFP were sequentially stained with antibodies that label pre- and post-synaptic proteins for multiplex fluorescence imaging. Example labeling in a single field of view with antibodies against the pre-synaptic marker Synapsin 1 (white), pre-synaptic glutamatergic marker VGLUT2 (magenta), pre-synaptic GABAergic marker Vgat (yellow), post-synaptic glutamatergic marker PSD95 (cyan), and post-synaptic GABAergic marker Gephyrin (red). D) Enlarged images of the inset in panel C demonstrating colocalization in Synapsin-1-expressing YFP-labelled *Sst+* terminals (*left*) of pre-synaptic proteins necessary for release of GABA (Vgat) and glutamate (Vglut2) (*top*) and the post-synaptic proteins for scaffolding of GABA (Gephyrin) and glutamate (PSD95) ionotropic receptors (*bottom*). E) Z-scored enrichment of antibody puncta within YFP+ boutons relative to that expected at random. Colors indicate data from the same image stack. Dashed lines represent ± 5 Z-scores. Total number of YFP labeled terminals=8493, 4 stacks from 3 animals. F) Average cross-correlations of Z-scored fluorescence intensities for all pairs of antibodies (n=4 stacks from 3 animals). G) Average co-variances of Z-scored fluorescence intensities for all pairs of antibodies within the YFP-labelled EP *Sst+* terminals.

Individual EP *Sst+* neurons express genes necessary for both glutamatergic and GABAergic transmission (Root et al., 2018; Shabel et al., 2014). To examine if individual synaptic boutons from these neurons in LHb express the proteins necessary for synaptic release of both glutamate and GABA, we used array tomography (Micheva and Smith, 2007). Cre-dependent expression of synaptophysin-YFP induced by AAV injection (AAV-DIO-Syn-YFP) into EP labeled *Sst+* presynaptic terminals in LHb. Serial sections were immunolabeled for YFP, Vglut2, VgatT, PSD95, and Gephyrin (Figure 1C-D). As expected, YFP was found in EP *Sst+* pre-synaptic terminals and colocalized with pre-synaptic protein marker Synapsin-1 (Figure 1D).

We hypothesized that if glutamate and GABA are released from the same pre-synaptic terminals, then the vesicular machinery for glutamate and GABA packaging (Vglut2 and Vgat, respectively) should co-localize. The relationships between the distributions of immunolabeled proteins were analyzed by two methods (Granger et al., 2020). First, individual boutons were identified and their boundaries determined from the YFP signal. Similarly, individual immunolabeled puncta for each antibody were identified and the centroid of fluorescence of each punctum was calculated (Figure 1D; Supplemental Figure 1A). To determine if specific antigens are preferentially localized in the YFP-defined boutons, we measured the fraction of YFP-positive pixels containing the centroid of an antibody punctum and compared it to that expected by chance (1000 randomizations of centroid locations) (Figure 1E). Synapsin immunopuncta were found within the YFP+ regions far more often than expected by chance (Figure 1E; Supplemental Figure 1B). Similarly, Vgat and Vglut2 immunolabeling often overlapped (Figure 1D) and puncta for both proteins were found in YFP-labeled terminals far-above chance (Figure 1E; Supplemental Figure 1C). In addition, we examined the overlap of YFP+ terminals with post-synaptic scaffolding proteins associated with glutamate (PSD95) and GABA (Gephyrin) receptors. We found strong non-random expression of Gephyrin overlapping with YFP+ boutons and weaker, but still above-chance, expression of PSD95 (Figure 1D-E; Supplemental Figure 1C).

For the second method of analysis, we avoided identifying individual immunopuncta and instead analyzed the cross-correlation and covariances of fluorescence intensities after normalizing each fluorescent channel independently to mean 0 and variance 1. Analysis of cross-correlations across each set of tissue images (Figure 1F) is dominated by immunolabeling outside of the YFP+ boutons, which cover on average only ∼0.3% of the image pixels (0.1-0.6% in 4 tissue stacks, 3 animals). Whole-image analysis revealed weak cross-correlations across all antibody channels (mean across samples: 0.003-0.294; individual samples: 0.0007-0.423), peaking at mean image displacement of 0. To focus analysis on the *Sst+* presynaptic terminals, we restricted analysis to the image areas within YFP-labeled terminals (Figure 1G). The Vgat-Vglut2 signal intensities had high positive covariance within the boundaries of YFP+ presynaptic terminals, indicating that glutamatergic and GABAergic vesicular transporters overlap in boutons of EP *Sst+* axons. Similarly analyzed Vgat-Gephyrin signals had high positive covariance, consistent with overlap of inhibitory pre- and post-synaptic densities for GABAergic terminals (Figure 1F-G). The signal from the PSD95 antibody did not exhibit positive covariance with any of the other antibodies, possibly due its low enrichment within the YFP+ boutons (Figure 1E, G) (Granger et al., 2020; Saunders et al., 2015).

Thus, individual EP *Sst+* presynaptic boutons in the LHb have the molecular machinery necessary to release both glutamate and GABA and colocalize with scaffolding proteins associated with GABA receptors. This indicates that individual boutons likely contain both transporters. However, due to the small size of synaptic vesicles compared to primary and secondary antibody complexes as well as to the limits imposed by the imaging resolution, these results cannot determine if glutamate and GABA vesicular transporters are found on the same vesicles.

### Statistical features of synaptic currents generated by two models of glutamate/GABA co-release

We considered two models that have been previously proposed regarding the mechanism of glutamate/GABA co-release in LHb: one in which the two neurotransmitters are packaged in separate vesicles but are released from the same terminal (Root et al., 2018) (termed the independent release model) and the other in which the two neurotransmitters are packaged in the same vesicles (Shabel et al., 2014) (termed the co-packaging release model) (Figure 2A). Under both scenarios the average PSCs produced by release from co-transmitting synapses, generated either by stimulating a single bouton many times or by pooling signals across many boutons, can appear identical. However, trial-by-trial analyses of synaptic currents resulting from stimulation of individual co-transmitting synapses differ in each model when vesicle release is stochastic (i.e. release probability, p_r_, is <1) (Figure 2A). Furthermore, in the independent release model, the maximum (i_max_) and minimum (i_min_) amplitudes are uncorrelated whereas in the co-packaging model, the amplitudes exhibit strong within-trial correlation (Figure 2B).

**Figure 2.**
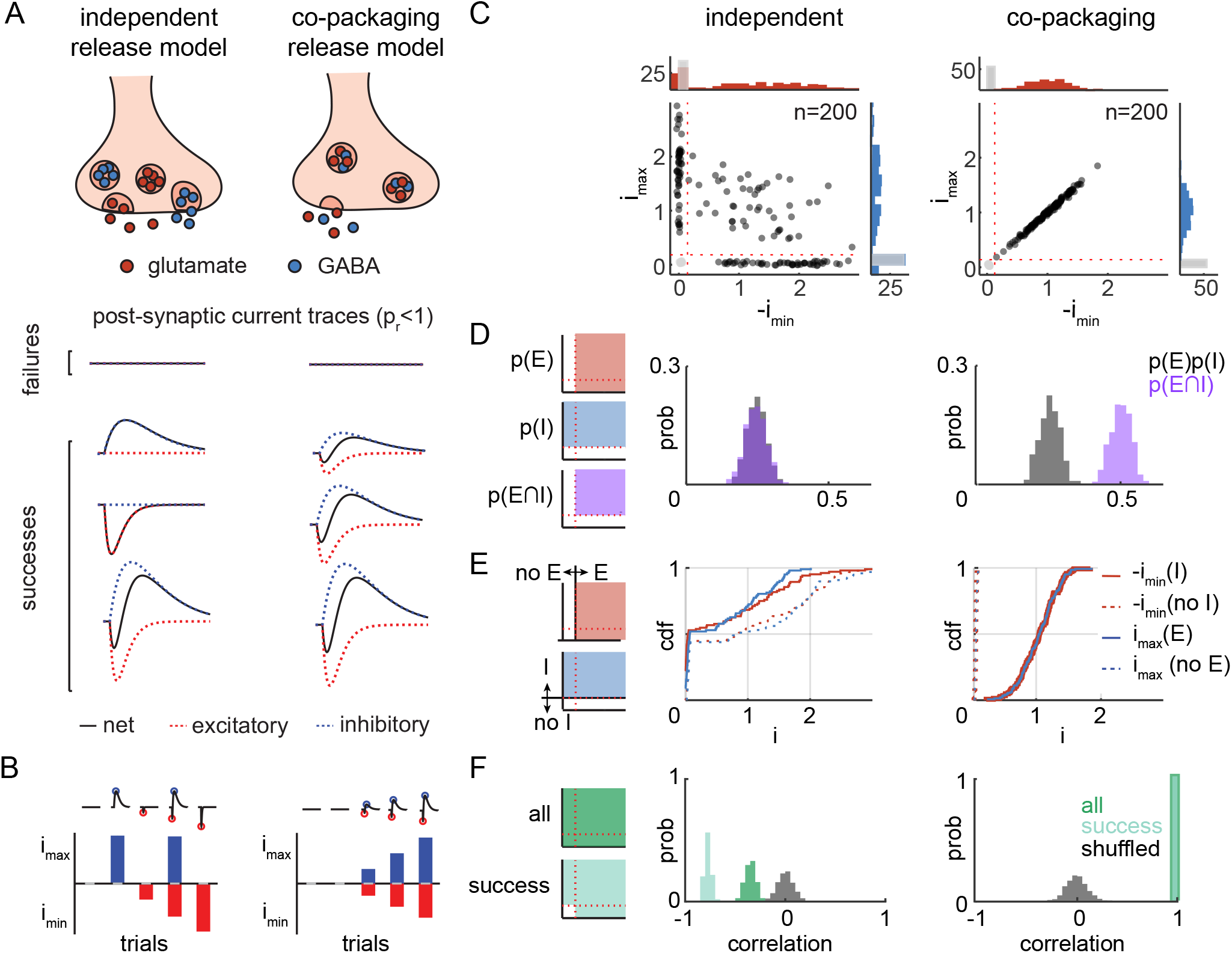
Statistical features of synaptic currents predicted by two models of glutamate and GABA co-release. A) *top*, Schematic showing two potential modes of glutamate and GABA co-release from individual synaptic terminals in which each class of vesicle is released independently (*left*) or the two neurotransmitters are co-packaged in the same vesicle and thus always released together (*right*). *bottom*, PSCs predicted by the independent (*left*) and co-packing (*right*) models at low synaptic release probability (p_r_). The independent model predicts that, following each stimulation of a single presynaptic bouton and successful vesicle release, the PSC can be excitatory, inhibitory, or biphasic. In contrast, the co-packaging model predicts that each PSC will be biphasic. In both models, failures of release can also occur. B) The maximum and minimum PSC amplitudes for the example trials in panel (A) for the independent (*left*) and co-packaging (*right*) release models. C) Scatterplots of the maximum and minimum amplitudes of 200 PSCs generated by simulations of independent (p_r_ = 0.5, *left*) and co-packaging (p_r_ = 0.75, *right*) release models with the same rates of synaptic failures (0.25). Amplitudes are normalized to the average maximum (y-axis) and minimum (x-axis) amplitudes of success trials. Histograms (in counts) of the normalized maximum and minimum release amplitudes with successes of release are shown on the right (blue) and top (red) and failures of release in each are shown in grey. D) *left*, Schematic representations of the areas within the scatterplots used to count events and calculate the probabilities of detecting inhibitory (p(I)) or excitatory (p(E)) currents as well as of biphasic currents with both inhibitory and excitatory components (p(E∩I)). Two different trial types contribute to p(E) and p(I), whereas only one trial type contributes to p(E∩I). *center and right,* Analysis of the statistical independence of the probabilities of detecting inhibitory (p(I)) and excitatory (p(E)) PSCs for the two models was generated by comparing the observed probability of excitatory and inhibitory PSCs (p(E∩I), purple) to that expected by chance (p(E)p(I), gray). Results for independent (*center*) and co-packaging (*right*) release models are shown with parameter p_r_ = 0.5. The summary of results from 1000 simulations are shown. For the independent model (*center*) the histograms overlap, largely obscuring the gray. E) *left*, Schematics representations of the areas within the scatterplots used to determine presence or absence of excitatory (*top*) or inhibitory (*bottom*) PSC for each trial. *center* and *right,* Simulated cumulative distribution functions (cdf) of maximum PSC amplitudes (i_max_, blue) given the presence (i_max_(E), solid) or absence (i_max_(no E), dashed) of an excitatory current in the independent (*center*) and co-packaging (*right*) release models. Similar analyses were performed for the minimum PSC amplitudes (-i_min_, red) given the presence (-i_min_(I), solid) or absence (- i_min_(no I), dashed) of an inhibitory current. Simulation parameters are the same as those used in panel D. F) *left*, Schematics of the areas of the scatterplots that contain all (*top*) or success-only (*bottom*) trials. *center* and *right,* Analysis of the trial-by-trial correlation of -i_min_ and i_max_ of all trials (dark green), success-only trials (light green), and after shuffling trial number labels across all trials to break the paired relationships between -i_min_ and i_max_ (grey). Results for the independent (*center*) and co-packaging (*right*) release models are shown. Simulation parameters are the same as those used in panel D.

To determine the features that can distinguish the two models, we implemented a biophysical simulation of the PSCs generated by stochastic synaptic vesicle release under either the independent or the co-packaging model (see Methods). Scatter plots of the maximum and minimum amplitudes extracted from PSCs generated by simulation of the independent release model revealed a dispersed distribution with 4 clusters of different synaptic responses (Figure 2C). In contrast, in the co-packaging release model, we observed 2 clusters with one corresponding to failure trials and one extending in a diagonal band that contains all the successful trials, consistent with the within-trial correlation between maximum and minimum amplitudes (Figure 2B). Moreover, clear differences are predicted by the two models in the population-level distributions of PSC amplitude maxima and minima when trials are grouped by failure and success, with the latter including EPSC-only trials, IPSC-only trials, and trials with both EPSCs and IPSCs (Figure 2C, see histograms along the top and right of each panel). In the co-packaging model, a clear separation is seen between failure and success trial maximum and minimum amplitudes (Figure 2C, *right*); whereas, in the independent release model, the maximum and minimum amplitude histograms of success trials cover a broader range, overlapping with those of failure trials (Figure 2C, *left*).

We calculated three statistical features from the simulated datasets that quantify the qualitative differences described above. These features differ in the degree to which they rely on the ability to accurately detect the presence of an EPSC or an IPSC in each trial (i.e. to distinguish successes from failures). Below we use the maximum (i_max_) and minimum (i_min_) current during a defined window to refer to amplitudes of inhibitory and excitatory currents without judging if a release event has occurred (i.e. they may be due to noise). In contrast, we use IPSC and EPSC and their amplitudes to refer to the components of PSCs that were judged to be a success of GABA or glutamate release, respectively (i.e. the excitatory or inhibitory component rises out of the noise – see Methods).

First, we considered the probabilities of detecting PSCs with different components. This method determines the presence or absence of the EPSC and IPSC on each trial but does not consider amplitudes of the detected currents. The occurrence of two events (e.g. detecting an EPSC or an IPSC) are statistically independent if and only if the probability of the events occurring together, or the joint probability, is equal to the product of the probabilities of each occuring. We adopted this framework to test if the observed probabilities of occurrence of PSCs with EPSCs, IPSCs, or both are consistent with the results predicted by statistical independence. Thus, we tested if:

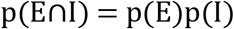

where p(E) is the measured probability of detecting an EPSC, p(I) is the measured probability of detecting an IPSC, and p(E∩I) is the measured probability of detecting a compound current in the same PSC (Figure 2D). As expected, only simulations of the independent release model generated a distribution of joint probabilities that matched the distribution of the products of the individual probabilities. Simulations of the co-packaging model produced a joint probability distribution shifted far right of the distribution predicted by independence probability theory (Figure 2D).

Second, we compared the histograms of PSC maximum and minimum amplitudes in trials grouped by the presence and absence of EPSCs and IPSCs (Figure 2E). This test examines if the minimum PSC amplitude distributions are the same whether or not an IPSC was detected in the trial (“I” or “no I” trials, respectively). The converse – the PSC maximum amplitude distributions for EPSC and no EPSC containing trials (“E” or “no E trials”, respectively) – was also examined. Thus, we calculated four cumulative distribution functions (cdfs).

In the independent model, the four cdfs rise sharply near zero amplitude, indicating that a failure of glutamate or GABA release does not predict the failure of release of the other transmitter (Figure 2E). Furthermore, the cdfs of PSC amplitudes from the “I” vs. “no I” trials show only small differences, consistent with the presence or absence of an IPSC having only small effects on the minimum amplitude. Similar observations are made for comparisons of the maximum amplitude cdfs of “E” vs. “no E” trials. In contrast, in the co-packaging model, the “no E” and the “no I” cdfs are each shifted far left relative to the “E” and “I” cdfs, respectively, consistent with the presence or absence of the one current fully predicting the presence or absence of the other current. Although this assay requires detecting the presence of either the EPSC or the IPSC on each trial, it is robust to some errors in the accuracy of detection. In fact, the requirement of judging the presence or absence of either component can be relaxed and the same analysis can be performed by simply dividing the PSC into those with, for example, large and small amplitude IPSCs and asking if this influences the distribution of EPSC amplitudes (Supplemental Figure 2A). The relaxed requirement still produces distinguishable differences between the two models, demonstrating that, even if signal-to-noise (SNR) of recordings is low, our statistical tests are robust.

Third, we examined the correlation coefficients across trials of the PSC minimum and maximum amplitudes (Figure 2F). Correlation analysis was performed separately for all trials and for success trials to account for possible analysis artifacts resulting from inclusion of noisy failure trials. In the independent release model, the distributions of the correlations between maximum and minimum PSC amplitudes are consistently negative when calculated for all trials and for success trials (Figure 2F). The negative correlation arises from the overlap of the EPSC and IPSC and reflects the differences observed in Figure 2C. Moreover, the success-trials correlation distribution is more negative compared to that for the all-trials correlation due to the algorithmic removal of the failure trials which, by definition, have noise-generated uncorrelated positive and negative deflections. In contrast, simulation of the co-packaging model produces strong positive correlations (essentially 1) for all-trials and for success-trials (Figure 2F). This high correlation results from (1) co-occurrence of successes and failures in EPSCs/IPSCs and (2) shared variance due to vesicle-to-vesicle size differences, which co-modulates the two opposing currents. In each case, null correlation distributions were computed by shuffling the maximum and minimum amplitudes across trials and, as expected, are centered at zero in both models (Figure 2F). This assay, when applied to all trials, does not require judging the presence or absence of either the EPSC or IPSC in each trial.

### DMD-based optogenetic stimulation to study glutamate/GABA co-release from EP *Sst+* axons

Previous studies of glutamate and GABA co-transmission at EP-LHb synapses have used wide-field optogenetic to evoke neurotransmitter co-release from many EP terminals while measuring compound PSCs in LHb neurons (Root et al., 2018; Shabel et al., 2014; Wallace et al., 2017). This produces essentially one response per postsynaptic neuron and obscures potential differences between individual terminals. A stable and repeatable method to target defined synapses is essential to statistically compare the experimental data with the predictions generated from computational simulations described above. We implemented a digital micromirror device (DMD)-based optogenetic stimulation approach to activate glutamate/GABA co-releasing EP *Sst+* axon terminals in the LHb (Figure 3A). The goal was to separately activate many different terminals as quickly and in as many trials as possible. This allowed us to measure the variance of stochastic neurotransmitter release across time and at different synapses. Variants of this approach (CRACM and sCRACM) were used to map connectivity and the spatial arrangement of synapses in cortical circuits (Petreanu et al., 2009, 2007). We adapted this approach to target small sets, ideally consisting of an individual (see below, Figure 4), presynaptic terminals.

**Figure 3.**
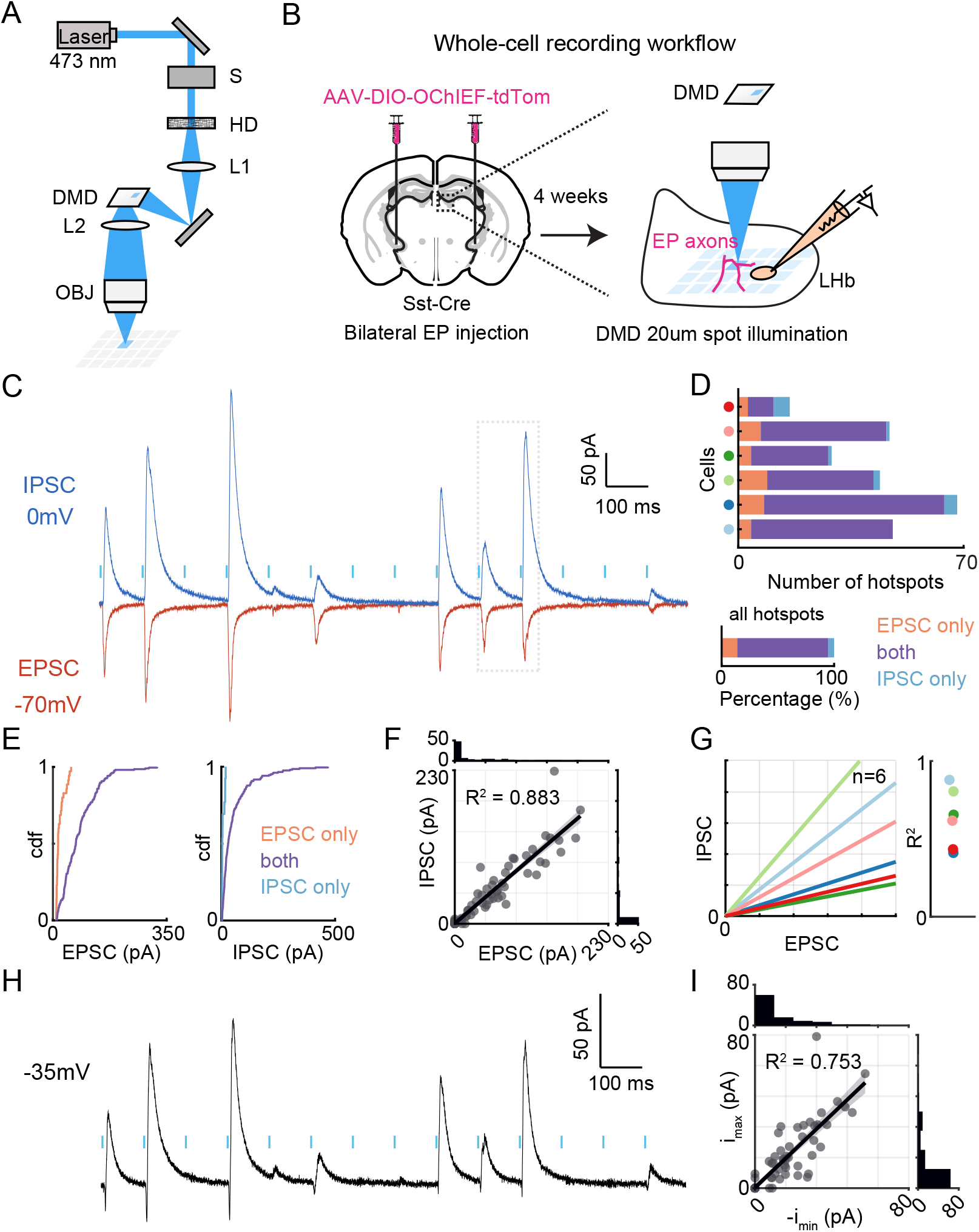
Optical approach to measure PSCs evoked by optogenetic stimulation of groups of EP Sst+ axons in the LHb. A) Schematic of the DMD-based minimal optogenetic stimulation (DMOS) platform. S: mechanical shutter; HD: holographic diffuser (10° diffusing angle); DMD: digital micromirror device; L1-2: lens; OBJ: objective lens. B) Schematic of the workflow showing injection of Cre-dependent AAV encoding the optogenetic activator OChIEF into the EP of *Sst-Cre* mice, followed by whole-cell recordings in acute-brain slices of LHb under the DMOS system. C) Optically-evoked average compound (due to high photo stimulation intensity) PSCs in an example LHb neuron. EPSCs and IPSCs were acquired while the cell was voltage-clamped at a holding potential (V_h_) of −70 mV (red) or 0 mV (dark blue), respectively. Light blue vertical bars show the timing of the laser pulses used for optogenetic stimulation with each pulse delivered to a different location in the slice. PSCs are the average of 5 trials. The dotted box encloses currents evoked at two stimulation spots that evoke EPSCs of similar size but IPSCs with widely differing amplitudes. D) The number of stimulation spots triggering PSCs (x-axis) in individual cells (*top,* y-axis) or across all cells (*bottom*) grouped by the presence of EPSCs only (orange), IPSCs only (blue), or both (purple) (n=6 cells/3 animals with 252 total active hotspots). E) Cumulative distribution functions comparing the EPSC and IPSC amplitude distributions in different classes of hotspots. *left*, EPSC-only hotspots have smaller EPSC amplitudes (orange) than do co-transmission hotspots (purple). *right*, IPSC-only hotspots have smaller IPSC amplitudes (blue) than do co-transmission hotspots (purple). Same dataset as in panel D. F) Scatterplot of all IPSC vs. EPSC peak amplitude pairs evoked at each photo-stimulated spot in an example LHb neuron. The IPSC/EPSC peak amplitude ratio is conserved across multiple sets of EP Sst+ axons synapsing onto the same post-synaptic target cell. The top and right histograms show the distributions of EPSC and IPSC amplitudes, respectively. Fitted line: *y* = 0.438 + 0.856*x*. G) Fitted IPSC/EPSC peak amplitude relationships for data from 6 LHb cells (*left*) and corresponding R^2^ values (*right*). Colors indicate cell identity matching as in panel D. H) Optically-evoked average biphasic, compound PSCs recorded at an intermediate holding potential, V_h_=-35 mV, in the same neuron as in panel C. Blue vertical bars show the timing of the laser pulses used for optogenetic stimulation with each pulse delivered to a different location under high photo stimulation intensity. PSCs are the average of 5 trials. I) Scatterplot of maximum and minimum current amplitude pairs in the PSCs recorded at −35 mV for the example trace shown in panel H. The top and right histograms show the distributions of minimum and maximum amplitudes of the PSCs, respectively. Fitted line: *y* = 0.316 + 0.955*x*.

**Figure 4.**
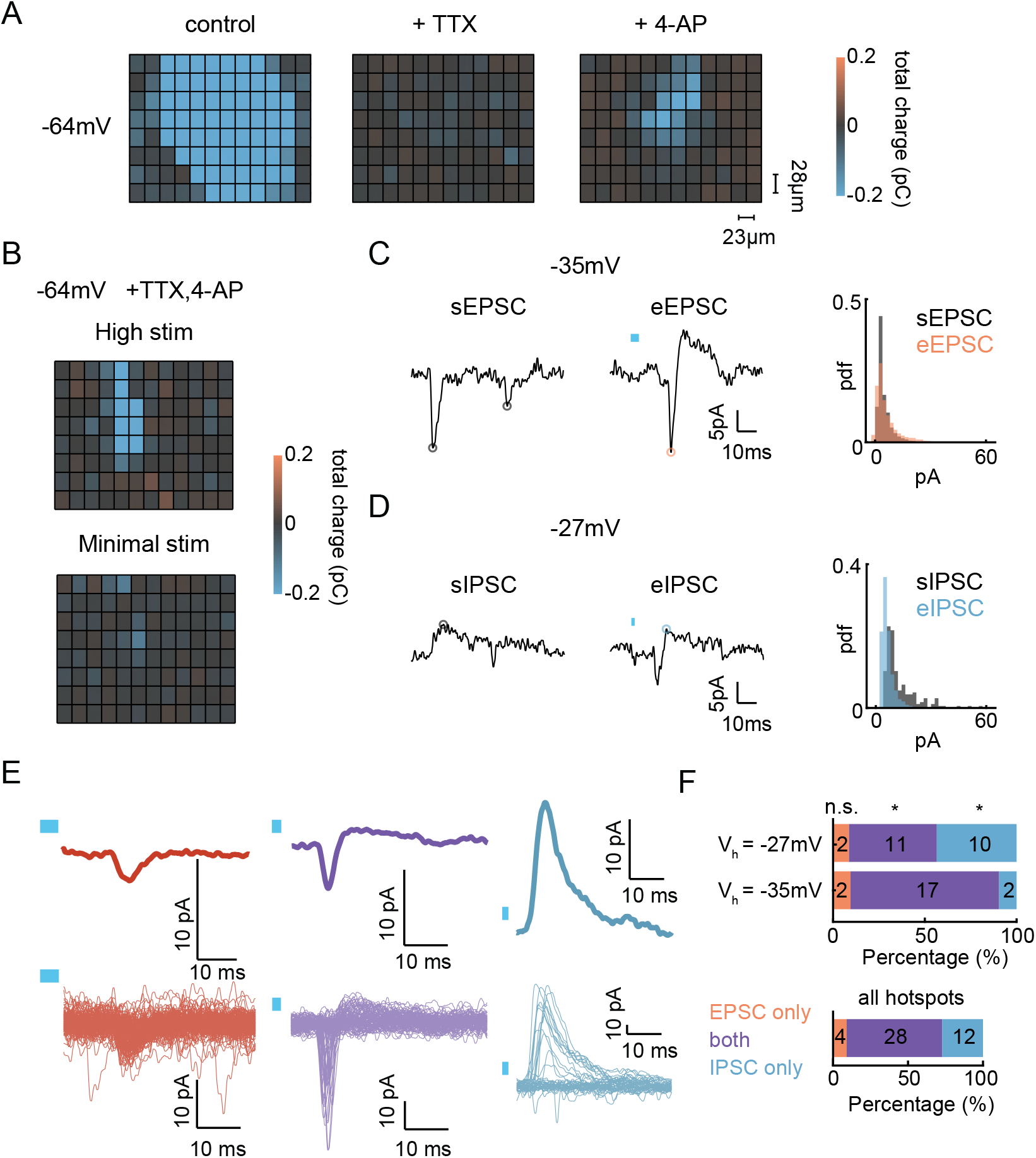
DMOS evokes unitary responses from EP *Sst+* axons in LHb. A) Example spatial heatmaps showing the effects of sequential addition of TTX and 4-AP on total charge of EPSCs (V_h_=-64mV) of all stimulation spots using DMOS under high photo-stimulation intensity. Total charge of PSC was measured in a 5-25ms time window after the onset of photo stimulation. Heatmap represents mean of 5 trials. The recorded cell was located approximately at the center of each heatmap. B) Example spatial heatmaps comparing total charge of EPSCs (V_h_=-64mV) of all stimulation spots using DMOS under high (*top*) and minimal (*bottom*) photo-stimulation intensity. Total charge of PSC was measured in a 5-25ms time window after the onset of photo stimulation. Heatmap represents mean of 5 trials. C) Examples of spontaneous EPSCs (sEPSCs) (*left*) and unitary evoked biphasic PSC (*middle*) with evoked EPSC (eEPSC) amplitude indicated. The PSC was evoked under minimal light stimulation in the same cell and holding potential at which the sEPSC was recorded. *right*, Histograms of peak amplitudes of sEPSCs (grey, median amplitude 95% CI = 3.37-3.50 pA, median frequency=8.9 Hz) pooled across all cells (14 cells, 9 animals) and eEPSCs measured across subset of the cells containing unitary evoked biphasic PSCs (orange, median amplitude 95% CI = 3.82-4.17 pA; 11 cells, 6 animals). Light blue boxes show the timing and duration of the laser pulses. Bin width of histogram is 2 pA. D) As in panel B for a spontaneous IPSC (sIPSC) and unitary evoked IPSC (eIPSC). The sIPSC (grey) had median amplitude 95% CI = 9.15-10.51 pA and frequency=0.2 Hz whereas the eIPSCs (blue) had median amplitude 95% CI =3.84-4.18 pA. E) Average (*top*) and individual (*bottom*) representative unitary PSCs recorded at an intermediate V_h_ and evoked by repetitive stimulation at three different spots that consistently evoked PSCs consisting of EPSCs only (red), IPSCs only (blue), or both (purple). Light blue boxes show the timing and duration of the laser pulses. F) The proportions of minimal stimulation spots that triggered PSCs in cells recorded at V_h_=-27 or −35 mV, as indicated (*top*), or across all cells (*bottom*, 14 cells from 9 animals with 44 total active minimally-evoked hotspots) grouped by the presence of EPSCs only (orange), IPSCs only (blue), or the presence of both EPSCs and IPSCs (purple). Asterisks indicate statistical significance of Fisher’s exact test comparison of the proportions of each group observed at −27 mV and −35 mV. Comparisons of the proportions of “both” and “IPSC only” groups across potentials reject the null hypothesis of no difference between the observed proportions at two holding potentials.

We first examined DMD-evoked responses at high laser powers that activate many synapses. We prepared acute coronal brain slices from LHb of *Sst-Cre* mice at least 4 weeks after bilateral stereotaxic injection of Cre-dependent AAV encoding the excitatory opsin OChIEF into the EP (Figure 3B, as in Figure 1A-B). The system enabled stimulation of 96 specific spatial targets, each a 23×28 µm box, in less than 10 seconds (Supplemental Figure 3A). LHb neurons were held in voltage-clamp mode at the reversal potentials of GABA_A_R (−70 mV) and AMPAR (0 mV) to isolate the excitatory and inhibitory PSCs, respectively (Figure 3C). In each neuron, a subset of the stimulation spots (252 of 576 spots, n=6 neurons; 16-68 of 96 spots per neuron) elicited synaptic currents. Over 80% of the spatial locations (204 of 252 spots) that evoked EPSCs also evoked IPSCs (Figure 3C-D). The amplitudes of EPSCs and IPSCs evoked at each spot were typically correlated in each cell but the IPSC/EPSC ratio (or slope of the correlation) varied from cell-to-cell (Figure 3F-G). The variability across different sets of synapses measured within the same cell was not due to differences in quality of voltage clamp (Supplemental Figure 3E-F). Nevertheless, there were spots that evoked EPSCs and IPSCs whose amplitude ratio was different than that of the other synapses onto the same cell (e.g. Figure 3C, dotted box), indicative of heterogeneity in the ratio of glutamatergic and GABAergic currents evoked by different synapses. The EPSC sizes of the “EPSC-only” spots and IPSC sizes of the “IPSC-spots” were significantly smaller than those of the “both” spots (Figure 3E), suggesting that the “EPSC-only” and “IPSC-only” sites might also contain IPSCs and EPSCs, respectively, that are below detection threshold.

Overall, these results are consistent with *Sst+* axons co-releasing and the post-synaptic cell being able to detect both transmitters. Control experiments to test the spatial specificity of DMD-based activation were performed, including examining the response pattern after moving the microscope objective by a known distance (Supplemental Figure 3B-C) and testing whether light leaks to nearby regions with increasing light intensity (Supplemental Figures 3D).

In a subset of cells, we examined if recordings at intermediate potentials (V_h_ = −27 or −35 mV) could be used to monitor the EPSC and IPSC simultaneously. We observed biphasic responses following photo-stimulation of the same spots at which isolated EPSCs and IPSCs were detected at each reversal potential (Figure 3H; Figure 3C). Amplitudes of the inward and outward peaks in the biphasic responses were highly correlated, consistent with the biphasic responses representing the summation of two opposite signed synaptic currents (Figure 3I). However, the range of inward and outward peak amplitudes was smaller compared to the measurements made at the reversal potentials due to (1) the mutual occlusion of the EPSC and IPSC and (2) reduction in driving force of synaptic currents (slope change from 0.856 to 0.955; R^2^ change from 0.88 to 0.75).

### Heterogeneity in unitary responses from EP *Sst+* co-releasing axons

In order to compare experimental data to the statistical models, it is necessary to study responses at individual synapses. We modified the conditions of spatially-specific DMD-based optogenetic activation to generate minimal responses and call this approach DMOS – <u>D</u>MD-based <u>m</u>inimal <u>o</u>ptogenetic <u>s</u>timulation. Whole-cell voltage-clamp recordings were performed in the presence of TTX and 4-AP to optogenetically activate pre-synaptic boutons without propagating action potentials (Figure 4A) (Petreanu et al., 2009). Furthermore, we tested a variety of stimulation intensities and spot sizes until we achieved EPSCs whose amplitudes were similar to those of miniature spontaneous EPSCs (mEPSCs) and that appeared stochastically trial-to-trial. Under these conditions, fewer of the stimulation spots evoked PSCs even when maximizing detection of inward currents (V_h_= −64 mV, Figure 4B).

Similarly, we performed whole-cell voltage-clamp recordings at an intermediate holding voltage, −35mV or −27mV, at which both EPSCs and IPSCs could be observed while minimally stimulating EP *Sst+* axons with TTX and 4-AP in bath. In each recording, we started with high intensity photo-stimulation and then lowered the light intensity until the DMOS-evoked PSC events became stochastic. These minimally-evoked PSCs were biphasic and the evoked EPSC (eEPSC) and IPSC (eIPSC) components had amplitudes similar to those of spontaneous EPSCs (sEPSC) and IPSCs (sIPSC) measured from all the recorded neurons, respectively (Figure 4C; Figure 4D) (the median and interquartile range (IQR) of amplitudes in pA for each current were: eEPSC: 4.0 (IQR 6.0); sEPSC: 3.4 (IQR 3.8); eIPSC: 5.4 (IQR 3.3); sIPSC: 9.7 (IQR 8.7)).

We found three types of evoked unitary PSCs (uPSCs) using the DMOS approach. In each terminal, across hundreds of trials, we either observed “EPSC-only” (left), “IPSC-only” (right), or “both” (middle) hotspots that revealed only EPSCs, only IPSCs, or both EPSC and IPSCs, respectively, on every success trial (Figure 4E). Overall, the majority (∼64%) of all uPSC hotspots (44 spots from 14 cells; 1-7 hotspots per cell with median of 2.5) exhibited both EPSCs and IPSCs, consistent with the co-packaging model (Figure 4F; Supplemental Figure 4C). This result was not affected by changing the detection threshold of EPSCs and IPSCs (Supplemental Figure 4B,D). We hypothesized that some of the “EPSC-only” and “IPSC-only” uPSCs result from occlusion rather than a true lack of IPSC and EPSC, due to reduction of ion channel driving forces at an intermediate holding voltage. Indeed, the relative proportion of “IPSC-only” hotspots increased to 34% (from 2/21 hotspots to 10/23 hotspots in 7 cells in each group, fisher’s test p = 0.0174) when the holding voltage was increased from −35 to −27 mV, suggesting that competition of opposing currents generated by EPSCs and IPSCs limits the detection of both signals (Figure 4F).

### Examples of unitary responses that support independent and co-packaging models

We investigated the three statistical features outlined above (Figure 2) for responses that showed “both” uPSCs. Note that the common failure modes of our analyses will artificially support a model of independent release of glutamate and GABA. For example, noise in the electrical recording that is incorrectly labeled as an evoked EPSC or IPSC, high spontaneous miniature spontaneous EPSC and IPSC (mEPSC/mIPSC) rates that result in spontaneous events being mislabeled as evoked, or activation of multiple terminals within a single stimulation spot will all tend to make co-packaging synapses appear as independently-releasing synapses.

Among DMOS-activated spots that generated biphasic PSCs, we found examples consistent with independent (Figure 5A-E) as well as co-packaging (Figure 5F-J) models based on the three statistical features described above. At sites consistent with independent release (e.g. Figure 5A), heterogeneous shapes of PSCs were observed across trials with minimum amplitude peaks (i_min_) typically preceding maximum amplitude peaks (i_max_) (Figure 5A), as expected for evoked EPSCs and IPSCs as opposed to noise. A scatter plot of i_min_ and i_max_ amplitudes (Figure 5B) revealed a dispersed pattern with a negative slope consistent with the independent model (compare with Figure 2C). Furthermore, a bootstrapped (n=10,000) probability distribution of detecting both i_min_ and i_max_ amplitudes in single trials was not different from that expected by chance (Figure 5C) and matched the probabilities generated when the natural paired relationship between the i_min_ and i_max_ was broken by shuffling one relative to the other (Figure 5C). Furthermore, we simulated (n=500 runs) the biophysical models of two different modes of co-release using the parameters (i.e. the number of trials, p(E), and p(I)) measured from the data collected in Figure5A-B with an assumption of high SNR. The distributions of the joint probability of detecting an EPSC and IPSC together matched that generated by the independent model (Figure 5C, *top*), and was clearly different from that generated by the co-packaging model (Figure 5C, *bottom*). Similarly, cdfs of the minimum amplitudes in trials with or without an IPSC showed no difference (Figure 5D), more consistent with the independent release model prediction (Figure 2E). The same was true for the maximum amplitude cdfs. Finally, bootstrapped (n=10,000) correlation distributions of maximum and minimum amplitude pairs were centered around zero for all-trials and slightly negative for success-trials (Figure 5E). Thus, this example of synaptic responses generated by DMOS-stimulation of one site 145 times are best described by a model of independent release of glutamate and GABA. It is unclear if this conclusion reflects true independent release and detection of glutamate and GABA at a single synapse, or potentially results from the confounds listed above such as the presence of both a glutamate-only and a GABA-only synapse in the illuminated site.

**Figure 5.**
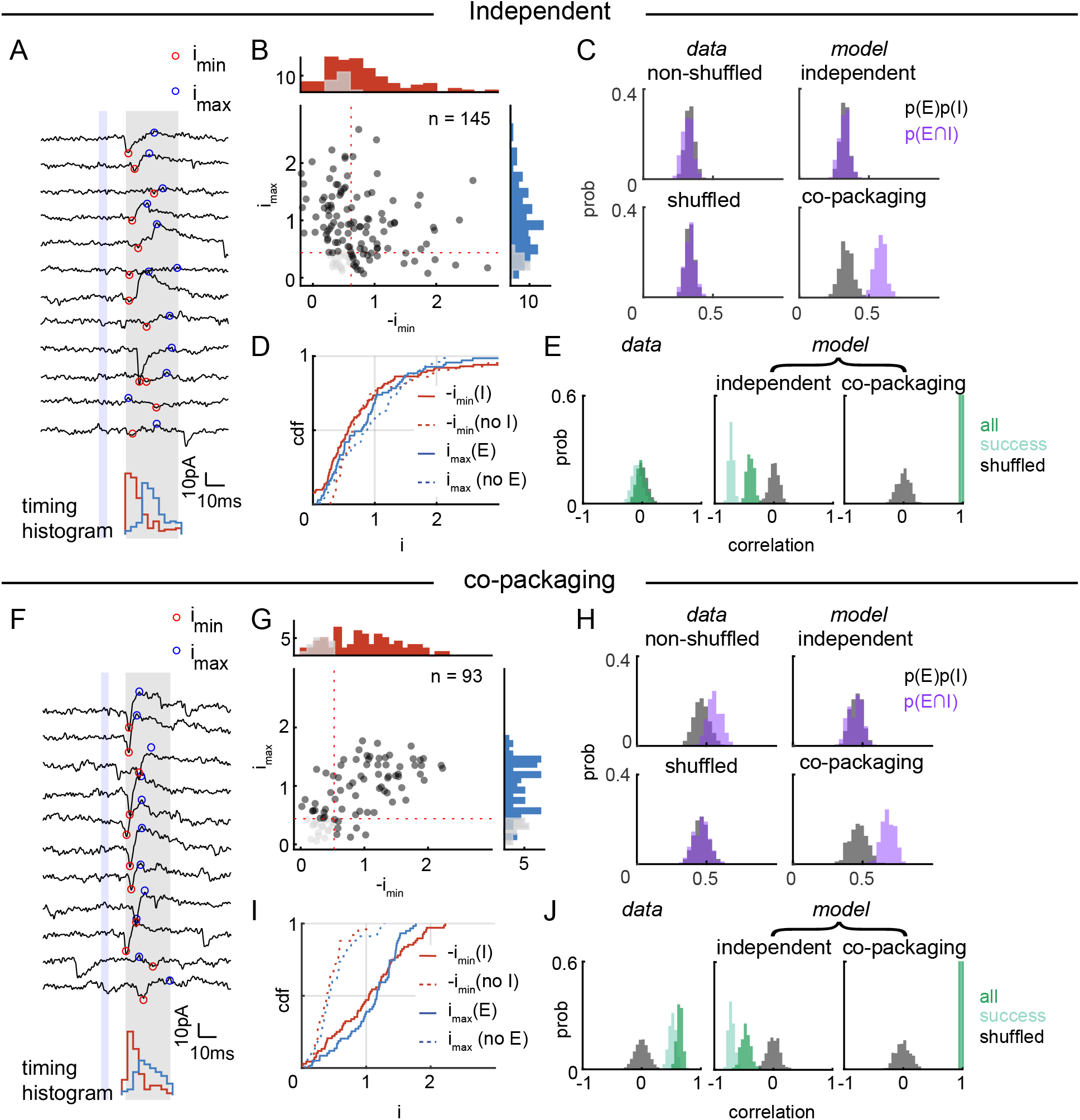
Unitary responses from glutamate and GABA co-releasing terminals. A) Optically-evoked PSCs from an example hotspot consistent with the independent release model. *top*, 12 example traces aligned to stimulus onset with blue shaded region indicating the duration of light stimulation delivered repeatedly to the same spot. The gray shaded region indicates the analysis window in which the maximum (blue dot) and minimum (red dot) amplitudes of the PSCs were extracted. *bottom*, Histogram of the times at which maximum (blue) and minimum (red) peaks were detected. B) Scatterplot of the maximum and minimum amplitudes of optically-evoked PSCs of the spot shown in panel A. Successes of release (either maximum or minimum amplitude exceeds the thresholds indicated by red dotted lines) trials are shown by black filled circles whereas failures of release are in gray. Histograms (in counts) of the maximum (*right*, blue) and minimum (*top*, red) release amplitudes are shown whereas amplitudes from failures trials are shown in grey. C) Analysis of the statistical independence of the probabilities of detecting evoked inhibitory (p(I)) and excitatory (p(E)) PSCs for the scatterplot shown in panel B determined by comparison of the observed probability of biphasic (excitatory and inhibitory) PSCs (p(E∩I), purple) to the probability expected by chance (p(E)*p(I), gray). *left,* Histograms of probabilities generated from boot strap analysis (10,000 repetitions) of actual data (non-shuffled, *top*) and shuffled data in which the pair-wise correspondence between maximum and minimum amplitude was lost (shuffled, *bottom*). The non-shuffled and shuffled datasets yield the same results, consistent with independent glutamate and GABA release at this site. *right*, Simulated histograms (500 repeats) of p(E)*p(I) and p(E∩I) generated by independent (*top*) and co-packaging (*bottom*) release models using synaptic parameters extracted from the data shown in panel B, showing that the data is most consistent with the independent release model. The areas within the scatterplots used to count events and calculate p(E), p(I) and p(E∩I) were set as in Fig 1. D) Cdf of maximum PSC amplitudes (i_max_, blue) given the presence (i_max_(E), solid) or absence (i_max_(no E), dashed) of an EPSC for the scatterplot shown in panel B. Similar analyses were performed for the evoked minimum PSC amplitudes (-i_min_, red) given the presence (-i_min_(I), solid) or absence (-i_min_(no I), dashed) of an IPSC. The areas within the scatterplots used to determine presence or absence of excitatory and inhibitory PSC for each trace are the same as in Fig 1. E) Analysis of the trial-by-trial correlation of -i_min_ and i_max_ across all trials (dark green), success- only trials (light green), and across all trials after shuffling trial number labels to break the natural relationship between -i_min_ and i_max_ (grey). Bootstrapped (10,000 repetitions) correlation coefficients for actual data (*left*) and correlation coefficient distributions from simulations (500 repetitions) of independent (*middle*) and co-packaging (*right*) release models are shown. The areas of the PSC amplitude scatterplots used to measure -i_min_ and i_max_ for all or success only trials are the same as in Fig 1. Analysis is of the data shown in (B) and using the same simulation parameters as in (C). F-J) As in panels A-E but for PSCs evoked at a hotspot with properties most consistent with the co-packaging model. In this case the IPSC and EPSC amplitudes are positively correlated (G and J); p(E∩I) is significantly greater than expected by chance (H); and the cdfs of -i_min_ and i_max_ depend on the presence or absence of on IPSC and EPSC, respectively (I).

At sites consistent with co-packaging, all successful event traces consisted of biphasic PSCs (Figure 5F). The scatter plot of the minimum and maximum amplitude pairs exhibited a positive correlation, with failures and success trials continuously spanning the diagonal axis of the distribution cloud (Figure 5G; compare with Figure 2C). The bootstrapped (n=10,000) probability distribution of detecting both an EPSC and IPSC was significantly greater (p < 1e-3) than the random distribution predicted by chance co-occurrence of an EPSC and IPSC (Figure 5H). The difference between the distributions disappeared when the EPSCs and IPSCs amplitudes were separately shuffled across trials. Furthermore, in agreement with the increased probability of detecting both EPSCs and IPSCs in single trials, this data was best fit by simulations of the co-packaging model rather than the independent model. In addition, cdfs of the minimum or maximum PSC amplitudes were well-separated when comparing across trials categorized by the absence vs. presence of an IPSC or EPSC, respectively (Figure 5I; compare with Figure 2E). Lastly, bootstrapped (n=10,000) trial-by-trial minimum and maximum amplitudes exhibited a large positive correlation for all trials and slightly smaller positive correlation for success trials (Figure 5J; compare with Figure 2F). The observed correlation of minimum and maximum amplitude pairs was not due to fluctuations of the stimulation intensity (Supplemental Figure 5A). Hence, our dataset contains example PSCs consistent with co-packaging of glutamate and GABA in the same vesicle, a conclusion that is difficult to arise artificially due to limitations of the methodology.

### Unitary responses of co-transmitting subtypes are consistent with the co-packaging release model

We performed the same analysis as above for each spot (n=28 from 11 cells) that exhibited DMOS-evoked biphasic PSCs. For each spot we performed the full analyses depicted in Figure 5A-E, including bootstrap-calculated distributions and comparison to simulation results generated by independent and co-packaging release models using the parameters tailored to each synapse. To quantify how much each statistical feature supported either model, a “model feature indicator” was parametrized to quantitatively capture the distribution differences described above (see Methods). In each case we compared the shift in the 50% value (i.e. the median) of two cdfs (Δcdf_0.5_), one presenting the data itself (or a distribution of bootstrapped data) and the other representing the equivalent cdf expected from chance observations of independent glutamate and GABA release (Figure 6A). This process resulted in 5 model feature indicators that summarize the deviation from random of each of the following: **1**, The joint probability of observing an EPSC and IPSC in the same trace; **2** and **3**, The influence of the presence or absence of an EPSC (2) or IPSC (3) on the amplitude of the IPSC (2) or EPSC (3); **4** and **5,** The correlation coefficients of i_min_ and i_max_ in each trial considering all trials (4) or success-only trials (5).

**Figure 6.**
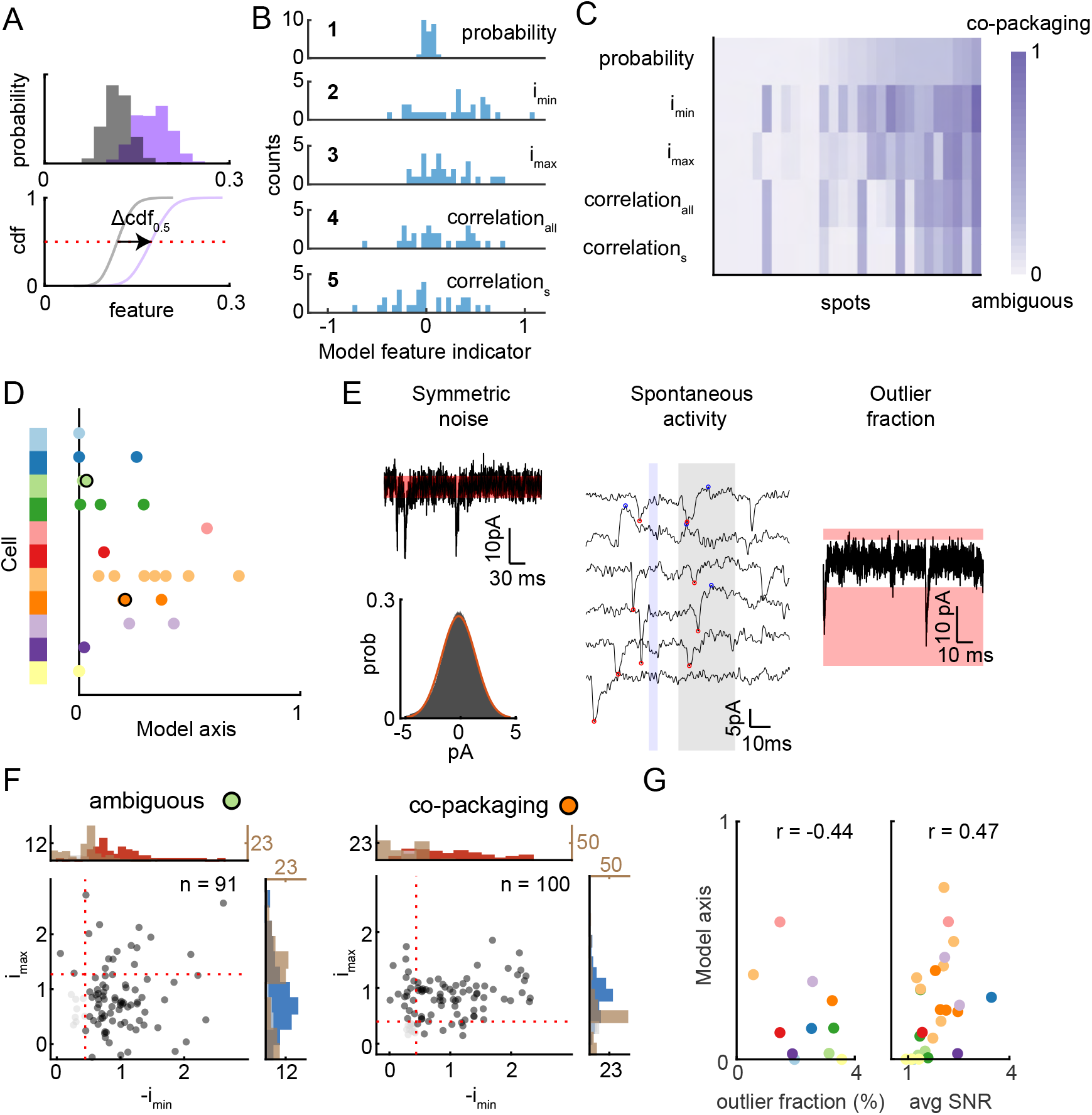
Statistical results of all unitary co-releasing terminals support the co-packaging model. A) Schematic illustrating the parametrization of a model feature indicator (Δcdf_0.5_) calculated by subtracting the medians of two cumulative distribution functions, one representing the distribution of p(E∩I) (purple) and the other that of p(E)*p(I) (grey). The difference between the x-values of the two distributions where cdf=0.5 indicates the direction and the strength of the relative shift of the feature distribution compared to that of the null. B) Histograms of the model feature indicators derived for the five statistical feature outputs. The data represents distributions of 28 DMD-evoked biphasic PSC spots from 11 cells (“both” group from Fig 3D). Bin width is 0.05. C) Heatmap of transformed model feature indicators from B (y-axis) across unitary spots exhibiting both EPSCs and IPSCs (x-axis). Color intensity represents increasing support for the co-packaging model. D) Distribution of average model feature indicators of all unitary co-releasing spots (n=28) based on 5 statistical feature outputs shown in C. Each dot represents an individual spot, with color indicating the identity of each cell. Increase model axis indicates greater support for the co-packaging model. Data collected from the black outlined spots are shown in detail in panel F. E) Schematic demonstrating three kinds of noise detected in the recordings. *left*, symmetric noise is the fluctuations around the baseline current that is fit by a gaussian function. *center*, minimum and maximum spontaneous PSC amplitudes can be detected due to spontaneous activity outside of the analysis window (grey shaded area) before the stimulus onset (blue shaded area). *right*, fraction of outlier current values (3x the scaled median absolute deviation away from the median of entire dataset) captures the contamination due to the frequency of large spontaneous synaptic current activity. F) Scatterplots of maximum and minimum amplitudes of an example ambiguous (*left*, green dot from D) and a co-packaging (*right*, orange dot from D) co-releasing hotspots. Histograms (in counts) of the evoked maximum (*right*, blue) and minimum (*top*, red) release amplitudes are shown whereas amplitudes from spontaneous activity during pre-stimulus baseline period are shown in brown (*right* and *top*). Note that the spontaneous activity histogram counts are scaled and shown in brown. G) Average model feature indicators for individual spots are correlated with parameters associated with signal-to-noise ratio, such as the fraction of outlier current values in a recording during the baseline period (*left*) and the average of EPSC and IPSC signal-to-noise ratio (SNR) (*right*). The SNR is calculated by dividing the evoked signal amplitude by twice the standard deviation of symmetric noise. Colors indicate cells identities, as in panel D. The Pearson correlation coefficient for each relationship is shown at the top of each plot.

Extremes values (i.e. near −1 or 1 except for the IPSC/EPSC joint occurrence probability feature which ranges between 0 and 1) of parameters indicated that categorization is strongly fit by either the co-packaging or independent release model. In contrast, values closer to zero reflected that the categorization was uncertain (Figure 6B). Unfortunately, the source of variations observed between −1 and 0 (i.e. those strongly fit by the independent model) is elusive as experimental errors can make co-packaging sites appear independent (see above and Discussion). As our study was designed to test if any synaptic responses were statistically compatible with co-packaging of glutamate and GABA, the model feature indicators were transformed to range from 0 (ambiguous or consistent with independent model release) to 1 (high confidence for co-packaging model) on the model axis (see Methods). The transformed model feature indicator heatmap of all sites revealed column-like structure (Figure 6C), indicating that the five statistical features captured in the model feature indicators are consistent as a group in their degree of support for the co-packaging release model (Supplemental Figure 6A). Using this metric, 22 of 28 sites had feature average greater than 0 (mean = 0.253, range 0.0057 ∼ 0.722) (Figure 6D).

As described above, the failure mode of our analyses is to favor the independent release model and false evidence for this model can result from high current noise, high spontaneous mEPSC/mIPSC rates, or the presence of multiple release sites in a single DMOS activated spot. To systematically investigate how these factors contribute to our results, we considered three noise metrics (Figure 6E). The quality of recording was captured by measuring the variance of the baseline current estimated from a gaussian fit. In addition, the impact of spontaneous mEPSC/mIPSC rates on the scatter distribution of observed trial-by-trial maximum and minimum amplitudes was measured. Lastly, we quantified the varying level of receptor saturation or kinetics across cells reflected in the spontaneous synaptic activity. Qualitative comparison between an example of an ambiguous site and a strongly supported co-packaging site (Figure 6F) revealed two major differences: (1) the separation between amplitudes of the spontaneous activity and those of minimum and maximum evoked currents; and (2) the SNR of evoked EPSC and IPSC amplitudes, which was calculated by comparing evoked EPSC/IPSC amplitudes to the EPSC/IPSC detection threshold limited by the baseline current noise.

At a population level, there was an inverse relationship between the degree of support for the co-packing model and the rate of spontaneous EPSCs/IPSCs, as judged by the fraction of outlier (i.e. 3x scaled median absolute deviation (MAD)) values in the holding current when no stimulus was delivered (Figure 6G; Pearson correlation coefficient = −0.44; p = 0.175). Furthermore, the average SNR of evoked currents was positively correlated (Pearson correlation coefficient = 0.47; p = 0.0112) with model feature indicator. Therefore, the sites with the best recording quality (low noise and low spontaneous synaptic events) had greater support for the co-packaging release model. This suggests that confounds of recording conditions may underlie the existence of sites that support the independent model or were ambiguous, such that most, if not all, co-transmitting sites might reflect synapses at glutamate and GABA are co-packaged.

### Pharmacological perturbation reveals co-packaging of glutamate and GABA in individual vesicles

A strong test of the co-packaging model is to examine if the correlations between glutamatergic and GABAergic currents (either their amplitude or simply their presence and absence) remain when probability of release is lowered. If both transmitters are in the same vesicle, then the co-occurrence of evoked inward and outward currents should persist when probability of release lowered. In contrast, if release of each transmitter is independent, then a 2-fold reduction of release probability should reduce the probability of biphasic currents 4-fold. Equivalently, if release is independent and lowered 2-fold, the detection rate of an IPSC when in a trial in which an EPSC is detected should fall 2-fold.

Serotonin reduces the probability of glutamate and GABA release from the EP axons in the LHb (Shabel et al., 2014, 2012) but it is unknown if serotonin has a similar effect on EP *Sst+* axons or equally on glutamatergic and GABAergic transmission. We examined the effect of serotonin (5-HT) on PSCs in LHb neurons resulting from activating groups of EP *Sst+* synapses (Figure 7A). We delivered an optogenetic ring stimulation using the DMD to evoke neurotransmitter release mediated by propagating action potentials and thus avoid direct activation of terminals synapsing onto the recorded neuron. This elicited composite excitatory and inhibitory PSCs in all cells (Figure 7B) (EPSC median (IQR) = 371.7 pA (385.7 pA); IPSC median (IQR) = 413.4 pA (423.6 pA); n= 6 cells, 5 animals), which were blocked by TTX (1 μM) and not recovered by 4-AP (400 μM), consistent with being evoked by propagating action potentials. Bath application of 5-HT (1 μM) reduced inward and outward currents in most cells (5 out of 6 for EPSC; 6 out of 6 for IPSC; unpaired t-test 5% significance level) (mean reduction: 19.6 ± 5.39% (EPSC), 40.9 ± 3.08% (IPSC); Supplemental Figure 7A), consistent with 5-HT mediated reduction of both glutamatergic and GABAergic release from EP *Sst+* axons in the LHb. These reductions in compound current amplitudes reflect the pooled effects of 5-HT on glutamate-only, GABA-only, and glutamate/GABA co-transmitting synapses (as in Figure 3).

**Figure 7.**
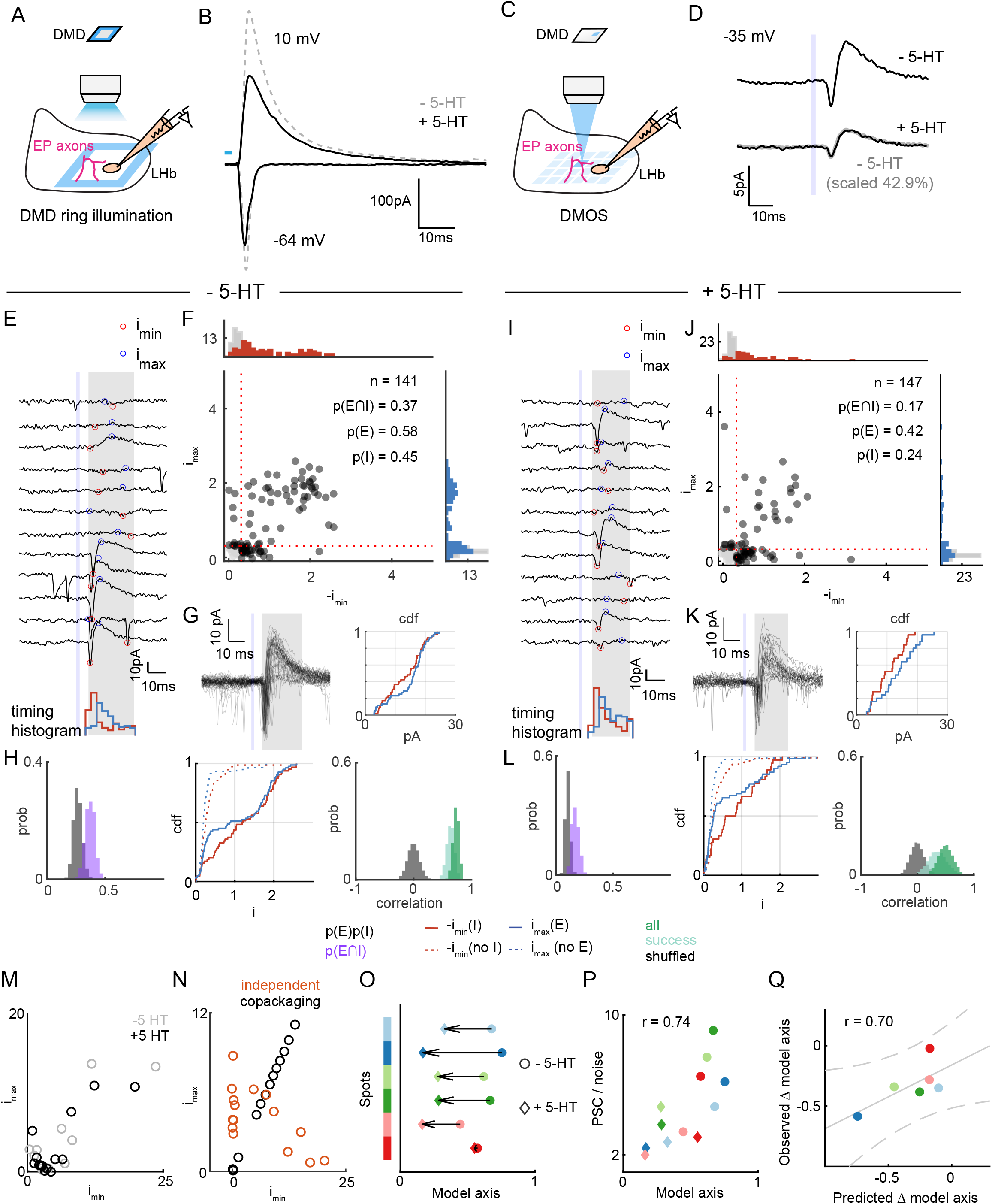
5-HT reduces probability of release of glutamate and GABA while maintaining their co-packaging. A) Schematic of 5-HT application experiment with DMD ring optogenetic stimulation. A LHb neuron was voltage clamped at −64 mV and 10 mV while the EP *Sst+* axons were optogenetically stimulated to generate propagating axon potentials, resulting in glutamate and GABA co-release. B) Example EPSC (V_h_=-64 mV) and IPSC (V_h_=10 mV) evoked by optogenetic activation of EP *Sst*+ axons using DMD ring photo-stimulation and recorded in a LHb neuron before (gray dashed) and after (black) bath application of 5-HT (1 µM). The blue box shows the timing and duration of the laser pulse. C) Schematic of 5-HT application experiment using DMOS to activate individual pre-synaptic boutons. D) Average biphasic PSC recorded from a LHb neuron (V_h_=-35 mV, black line)) following optogenetic activation of an EP *Sst*+ bouton using DMOS before (*top*, n=141 trials) and after bath application of 5-HT (250 nM, *bottom,* n=147 trials). 5-HT proportionally reduced the average biphasic response – the average biphasic response before 5-HT application is shown scaled and overlaid (grey) on the bottom. The shaded blue box shows the timing and duration of the laser pulse at minimal intensity. E) Optically-evoked PSCs from an example hotspot consistent with the co-packaging model. *top*, 12 example traces aligned to stimulus onset with blue shaded region indicating the duration of light stimulation delivered repeatedly to the same spot. The gray shaded region indicates the analysis window in which the maximum (blue dot) and minimum (red dot) amplitudes of the PSCs were extracted. *bottom*, Histogram of the times at which maximum (blue) and minimum (red) peaks were detected. F) Scatterplot of the maximum and minimum amplitudes of optically-evoked PSCs for the spot shown in panel E. Successes of release (either maximum or minimum amplitude exceeds the thresholds indicated by red dotted lines) trials are shown by black filled circles whereas failures of release are in gray. Histograms (in counts) of the maximum (*right*, blue) and minimum (*top*, red) release amplitudes are shown whereas amplitudes from failures trials are shown in grey. The probabilities of detecting an EPSC, IPSC, and both are shown in the inset. G) *left*, optically-evoked PSCs (V_h_=-35 mV) showing successes of both neurotransmitter releases. *right,* cumulative distribution function of the maximum (blue) and minimum amplitudes (red) of these trials. H) Analysis of statistical features shown: *left,* statistical independence of the probabilities of detecting evoked inhibitory (p(I)) and excitatory (p(E)) PSCs for the scatterplot shown in panel F determined by comparison of the observed probability of biphasic (excitatory and inhibitory) PSCs (p(E∩I), purple) to the probability expected by chance (p(E)*p(I), gray). *middle,* cdf of maximum PSC amplitudes (i_max_, blue) given the presence (i_max_(E), solid) or absence (i_max_(no E), dashed) of an excitatory current and vice versa for the scatterplot shown in panel F. *right,* trial-by-trial correlation of -i_min_ and i_max_ across all trials (dark green), success-only trials (light green), and across all trials after shuffling trial number labels to break the natural relationship between -i_min_ and i_max_ (grey). I-L) As in panels E-H after 5-HT (250 nM) bath application for the same hotspot. M) 5-HT effects on subset distributions of maximum and minimum amplitudes for the same site shown in E-L, without sorting trials by success and failures. Dots show the amplitudes of the average trace for different subsets of the dataset before (grey) and after (black) 5-HT application. N) As in M for showing the distributions predicted by independent (orange) and co-packaging (black) model simulation results. O) The effect of 5-HT on the distribution of average model feature indicators of unitary co-releasing spots consistent with co-packaging model (n=6) based on 5 statistical feature outputs shown in Figure 6C. Colors indicate spot identity. Arrows indicate the direction of the model feature indicator change due to 5-HT application. Circles indicate before 5-HT whereas diamonds indicate after 5-HT bath application condition. P) Average ratio between PSC and noise of individual spots versus average model feature indicators. Average PSC/noise ratio is calculated by dividing average minimum and maximum amplitude of all trials by the standard deviation of the baseline noise of a given cell. Colors and markers are as in panel O. Pearson correlation coefficient is shown at the top. Q) Comparison of observed model axis change due to 5-HT and that predicted by model simulation with updated release probability, changes in noise fluctuations, and changes in the evoked amplitude of “both” trials. Colors are as in panel O. Linear regression fit is shown in solid grey and the estimate of 95% prediction interval is shown in dashed grey. Pearson correlation coefficient is shown at the top.

To test whether 5-HT modulates biphasic PSCs resulting from activation of individual EP *Sst+* terminals, we examined the effects of 5-HT on DMOS-evoked hotspots with characteristics consistent with the co-packaging model (Figure 7C). Application of a low concentration of 5-HT (0.25 μM) reduced both inward and outward current amplitudes of average biphasic PSCs (Figure 7D, Supplemental Figure 7B; mean reduction: 45.4 ± 8.08% (-i_min_); 63.2 ± 4.18% (i_max_); mean number of trials: 123 ± 8.91; n= 6 spots, 6 cells, 6 animals). Trial-by-trial analysis indicated that, prior to 5-HT application, successful release trials consisted of biphasic PSCs with inward current followed by outward current (Figure 7E), consistent with earlier results (Figure 5F). Moreover, distributions of trial-by-trial maximum and minimum amplitude peaks and the three statistical features were consistent with those predicted by the co-packaging model (Figure 7F,H; Figure 5F-J). 5-HT reduced probability of success trials (mean reduction: 20.3 ± 4.94%, unpaired t-test p < 1e-3), probability of detecting EPSC (mean reduction: 22.9 ± 5.22%, p < 1e-3), IPSC (mean reduction: 30.3 ± 5.15%, p < 1e-3), and both (mean reduction: 32.9 ± 3.45%, p < 1e-3) (Figure 7J,L; Supplemental Figure 7C). Thus, 5-HT reduces both GABA and glutamate release from individual terminals that appear to package both transmitters in individual vesicles.

The distributions of i_min_ and i_max_ amplitudes spanned similar ranges before and after 5-HT bath application (Figure 7F,J). Waveforms and the cdfs of the i_min_ and i_max_ amplitudes of the “both” success trials were comparable (Figure 7G,K) and the bootstrapped Kolmogorov-Smirnov (K-S) tests (10,000 times) indicated no significant difference between the two groups (mean i_min_: 13.5 pA (before), 10.4 pA (after); mean i_max_: 14.8 pA (before), 13.36 pA (after); number of trials: 52 (before), 25 (after); p = 0.4804 (i_min_), p = 0.6891 (i_max_); Supplemental Table 1). In the same dataset, cdfs of the i_min_ and i_max_ amplitudes of all success trials were not significantly different (mean i_min_: 9.30 pA (before), 7.02 pA (after); mean i_max_: 9.55 pA (before), 6.54 pA (after); number of trials: 94 (before), 72 (after); Supplemental Table 1). Only two out of six cells had significantly different cdfs of i_min_ amplitudes and only one out of six cells had a significant i_max_ cdf difference (Supplemental Table 1) in the “both” success trials. Thus, the major effect of 5-HT on DMOS-evoked uPSCs is to reduce probability of release; however, 5-HT may have additional effects on post-synaptic receptor opening (i.e. synaptic potency).

To specifically test if the correlation between glutamate and GABA receptor currents was maintained after 5-HT application as predicted for the co-packaging model, we developed an alternative test that uses paired data from the basal and drug condition but does not require sorting trials into successes and failures. We compared the distribution of i_min_ and i_max_ amplitudes in trials sorted and binned by i_min_ amplitude – i.e., the 5 trials with largest i_min_ in group 1, the next 5 largest in group 2, etc… A positive correlation of the binned distributions of i_min_ and i_max_ confirmed that these sites were consistent with the co-packaging model (Pearson correlation coefficient = 0.893 (before), 0.817 (after) ; p < 0.001 (before and after)) (Figure7M; Supplemental Figure7D). Co-packaging vs. independent release models make different predictions of the effect of 5-HT on this relationship. In the former, assuming no change in synaptic potency, the range of the data and slope of the relationship showed remain unchanged; indeed, this was the effect observed in the example site (Figure 7M). If there is an additional change in synaptic potency, the relationship should scale along the diagonal whereas, if the effects are differential on glutamate and GABA receptors, the relationship should change slope. In contrast, in an independent release model in which the pre-5-HT consistency with co-packing arose by change, the relationship should be randomized after 5-HT or possibly reveal a negative correlation reflecting the mutual occlusion of excitatory and inhibitory synaptic currents (Figure7N).

Overall, we found that, after 5-HT application, the binned i_min_ vs. i_max_ distribution maintained the correlation slope in 3 out of 6 spots (Supplemental Figure7D1, D4, and D6). In the remaining the three spots, a correlation was maintained but the data shifted, consistent with larger effect on the i_max_ (i.e. IPSC amplitude) distribution (Supplemental Figure7D2, D3, and D5). Such effects could arise from a larger effect on potency of GABAergic vs. glutamatergic currents or reflect AMPA receptor saturation in the larger excitatory currents.

In addition, after 5-HT bath application, the three statistical features in all sites continued to support the co-packaging model (Figure 7H,L; Figure 7O; n= 6 out of 6 spots). In all cases the mean model indicator value continued to be positive and support co-packaging. Nevertheless, the mean model indicator decreased on average by −0.33 ± 0.07 (Supplemental Table 1), as expected from a reduction of SNR due to effects on synaptic potency or increases baseline noise and run down of synaptic currents that invariably occurs during long recordings. Indeed, model indicator values pooled from two conditions were strongly correlated with the ratio of the average PSC amplitude and current noise level of the individual spots (Pearson correlation coefficient = 0.74, p = 0.0063) (Figure 7P). Importantly, changes in the release probability, baseline noise, and PSC amplitude in “both” success trials accounted for the observed changes in model indicator value (Pearson correlation coefficient = 0.70, p = 0.12, norm of residuals of fit = 0.29) (Figure 7Q). These results demonstrate that 5-HT reduces release probability of both glutamate and GABA from EP *Sst+* inputs to the LHb and that terminals with features consistent with co-packaging continue to exhibit these features after reductions in probability of release.

## Discussion

Here we describe a novel experimental and statistical analysis approach to test distinct mechanistic models of neurotransmitter co-transmission. The approach is generally applicable to study synapses at which co-transmission is thought to occur and we apply it to examine glutamate/GABA co-transmission at EP *Sst+* terminals in LHb. We identify three statistical features that differentiate between computational models, one in which glutamate and GABA are released independently and another in which they are packaged in the same synaptic vesicle. Experimental data collected by activating individual pre-synaptic terminals reveal heterogeneity in neurotransmitter co-transmission. Nevertheless, we demonstrate examples of synapses that, when repetitively activated by minimal optogenetic stimulation, generate PSCs whose properties are consistent with co-packaging of glutamate and GABA and incompatible with independent release of each transmitter. Furthermore, pharmacological perturbations confirm that the statistical properties expected from co-packaged release of glutamate and GABA are preserved when release probability is lowered. Lastly, analysis of the contributions of synaptic noise and recording quality suggest that many synapses labeled as more consistent with independent release of glutamate and GABA, may actually reflect co-packaged release but with the expected correlations between glutamatergic and GABAergic currents obscured by noise. Thus, we conclude that EP *Sst+* neurons package both glutamate and GABA into the same vesicles and release these to activate correlated excitatory and inhibitory currents in LHb neurons. These findings have important implications for the plasticity mechanisms employed at this synapse, the relationship between activity in the EP and LHb, and maladaptive states known to induce plasticity in the circuit such as chronic stress, depression, and addiction.

### EP *Sst+* axons form glutamate/GABA co-releasing synapses in LHb

Here we exploited a *Sst-Cre* transgenic mouse to exclusively focus on glutamate/GABA co-releasing projections from EP to LHb. We found enrichment of the glutamate and GABA vesicular transporters, Vglut2 and Vgat, respectively, in EP *Sst+* terminals. High covariance of expression of these two pre-synaptic proteins agrees with analyses using immunogold electron microscopy that supports the conclusion that glutamate and GABA are released from EP and other terminals in the LHb (Root et al., 2018; Shabel et al., 2014). Curiously, we find that post-synaptic scaffolding protein Gephyrin, but not PSD95, is highly enriched near EP *Sst+* terminals despite the clear glutamatergic nature of these boutons (Li et al., 2011; Maroteaux and Mameli, 2012). This may indicate that, in contrast to glutamatergic terminals in cerebral cortex and hippocampus, an alternative MAGUK protein forms the core of these post-synaptic terminals. A positive correlation between Vglut2 expression and that of Synapsin-1 and PSD95 globally (i.e. in all terminals in LHb) (Figure 1G) indicates the existence of other molecularly distinct glutamatergic synapses in LHb (Barker et al., 2017; Hu et al., 2020; Knowland et al., 2017; Stamatakis et al., 2016).

The existence of glutamate and GABA co-packaging vesicles had been initially proposed following the observation of biphasic spontaneous miniature synaptic currents in LHb neurons (Shabel et al., 2014). However, the source of these biphasic miniature responses detected in LHb neurons were unknown since LHb receives projections that release glutamate and GABA from several brain regions (Barker et al., 2017; Stamatakis et al., 2016), including the ventral-tegmental areas (VTA) (Root et al., 2018, 2014). Based on our data, we propose that EP *Sst+* terminals are the source of synaptic vesicles that co-package glutamate and GABA. Interestingly, although the VTA also sends glutamate/GABA co-releasing axons to LHb, these are thought to release each transmitter from a separate pool of vesicles (Root et al., 2018).

Previous studies examined EP to LHb projections from the perspective of cellular physiology, anatomy, behavior, and disease models (Meye et al., 2016; Root et al., 2018; Shabel et al., 2014, 2012; Stephenson-Jones et al., 2016) using electrical stimulation or bulk channelrhodopsin activation of molecularly undefined, Vglut2+, or Vgat+ EP inputs in LHb. However, EP to LHb projections consist of two distinct neural populations that both normally express *Slc17a6* (encoding Vglut2) and hence express Cre in Vglut2-Cre (*Slc17a6-Cre)* mice. One population is *Sst+*, the glutamate and GABA co-releasing population studied here, and the other is Parvalbumin positive (*Pvalb+*) and purely glutamatergic (Wallace et al., 2017). Hence, these previous studies likely examine the structure and function of axons in LHb arising from both populations.

### Function of the EP, LHb, and co-release

LHb-projecting EP neurons, which include both *Pvalb+* and *Sst+* neurons, receive inputs from limbic-associated striosomes in the striatum (Wallace et al., 2017) and their firing rate is increased by aversive outcomes and decreased by rewarding outcomes (Hong and Hikosaka, 2008; Stephenson-Jones et al., 2016). Stimulation of all LHb-projecting EP neurons is aversive and impacts evaluation of action-outcome, thereby biasing future choices (Shabel et al., 2012; Stephenson-Jones et al., 2016), although it remains to be determined whether these populations contribute sufficiently to drive this effect (Lazaridis et al., 2019). Importantly, EP *Vgat+* neurons that project to LHb (putative *Sst+* neurons) preferentially target LHb neurons projecting midbrain GABAergic neurons (Meye et al., 2016), suggesting a function of EP *Sst+* neurons regulating the dopamine system. Since increased LHb activity can have aversive and reinforcing effects (Lammel et al., 2012; Proulx et al., 2014; Stamatakis and Stuber, 2012), the net ratio of glutamate and GABA released from EP *Sst+* terminals may determine the behavioral consequence resulting from modulation of these cells.

EP *Sst+* inputs transmit with similar ratio of glutamate/GABA currents to the same postsynaptic LHb neuron (Figure 3F) but it is unclear whether a pre- or post-synaptic mechanism underlies this phenomenon. Interestingly, the glutamate/GABA current ratio differs across different post-synaptic LHb neurons (Figure 3G), suggesting that each post-synaptic LHb neuron can integrate a unique combination of information from the same set of EP *Sst+* inputs by separately varying numbers of glutamate and GABA synaptic receptors. We speculate that glutamate and GABA co-transmission achieved by co-packaging in the same vesicles with post-synaptic variability in numbers of glutamate and GABA receptors allows each LHb neuron to use graded and signed synaptic weights assign to its inputs the combination of weights that best predicts an aversive outcome. Thus, negative weights are assigned to inputs whose activity coincides with or predicts a good outcome and positive weights are assigned to those associated with bad outcomes.

### Technical concerns involving study of glutamate/GABA co-release

The success of our analysis method depends on the SNR of the recording and the ability of the algorithm to detect glutamate or GABA-mediated currents with differing kinetics and amplitudes. The performance of the algorithms and the power of the models depend on the EPSC/IPSC transmission ratio and receptor kinetics and degrade with increasing spontaneous synaptic activity, baseline noise, electronic noise, and numbers of active terminals within an optogenetic stimulation spot. These factors tend to make co-packaging synapses appear as independent synapses. Indeed, our study finds that the likelihood of individual unitary response hotspots being categorized as co-packaging synapse is anticorrelated with level of spontaneous synaptic input firing level and correlated with average EPSC/IPSC SNR of the synapse (Figure 6E-G).

In this study, the ability to detect glutamate and GABA release depends on the expression of ionotropic receptors for each transmitter in the post-synaptic terminal associated with the stimulated bouton. Therefore, we are unable to state if synapses in which we observe only glutamate or only GABA mediated currents reflect terminals that release only one transmitter or post-synaptic terminals that are exposed to both transmitters but lack one of the receptor classes. Furthermore, given the small size of unitary synaptic currents and the ability of excitatory and inhibitory currents to occlude each other, in some glutamate-only or GABA-only spots it is possible that the missing current was simply hidden.

A possible source of error that could make independent sites appear as co-packaging sites is large variability in stimulation intensity that drives the correlation of amplitudes observed across trials. In this case the stimulation intensity would have to vary sufficiently to stochastically excite one or a small set of synapses that independently release glutamate and GABA, but do so with probability of release near 1. To test for this possibility, we measured the DMOS photo-stimulation intensity and demonstrated that trial-to-trial variations in stimulation intensity are small (<1%) and uncorrelated with the categorization of each trial as success or the amplitude of the EPSC and IPSC in a given trial (Supplemental Fig 5B).

### Serotonin modulation of glutamate and GABA co-releasing neurons

Application of serotonin reduces the amplitude of glutamatergic and GABAergic currents evoked in LHb neurons by stimulation of EP *Sst+* axons. This is consistent with previous findings that showed serotonin reduces the probabilities of glutamate and GABA release from EP (Shabel et al., 2014, 2012). We find that in synapses that co-package glutamate and GABA, the effects of 5HT are through largely mediated by a decrease in the probability of release of these vesicles with a potential additional effect on synaptic potency. 5HT receptor subtype 1B (5HTR_1B_) expressed in EP *Sst+* neurons likely mediates the presynaptic effect (Hwang and Chung, 2014; Wallace et al., 2017).

Serotonin signaling in LHb has been investigated in context of depression and its treatment. In animal models of depression, presynaptic changes have been described that shift the ratio of EP-to-LHb glutamatergic to GABAergic transmission, and this effect is reversed by treatment with selective serotonin reuptake inhibitor (SSRI)-type antidepressants (Shabel et al., 2014). Although our results suggest that the short-term effect of 5-HT is to inhibit release from *Sst+* inputs in LHb, longer-term additional effects of 5-HT on glutamate/GABA co-packaging vesicles remain unknown.

The interdisciplinary approach demonstrated here can be used to examine other co-transmitting synapses in the central nervous system to gain a richer understanding of all forms of neurotransmitter co-release.

## STAR Methods

### Mice

*Sst-Cre* (Jackson Labs #013044; MGI #4838416) homozygous and heterozygous mice (C57BL/6; 129 background) were bred with C57BL/6J mice. Both sexes of mice between 2-6 months in age were used. All animal care and experimental manipulations were performed in accordance with protocols approved by the Harvard Standing Committee on Animal Care following guidelines described in the US NIH *Guide for the Care and Use of Laboratory Animals*

### Viruses

To achieve specific expression of light-gated cation channel in the *Sst+* population in EP, we used a Cre-dependent adeno-associated virus (AAV) that encodes oChIEF, a variant of channelrhodopsin (Lin et al., 2009), driven by the EF1a promoter (AAV8-EF1a-DIO-oChIEF(E163A/T199C)-P2A-dTomato-WPRE-BGHpA). The plasmid was commercially obtained from Addgene (#51094) and the AAV was packaged by Boston Children’s Hospital Viral Core. For intracranial injections, the virus was diluted to a titer of ∼9 x 10^12^ gc/ml.

### Intracranial Virus Injections

Adult mice (>P50) were anesthetized with 2-3% isoflurane. Under the stereotaxic frame (David Kopf Instruments), the skull was exposed in aseptic conditions and the virus was injected bilaterally into the EP (coordinates: −1.0mm A/P, +/- 2.1mm M/L, and 4.2mm D/V, from bregma) through a pulled glass pipette at a rate of 50 nl/min with a UMP3 microsyringe pump (World Precision Instruments). 150 nl was infused per injection site. At least 4 weeks passed after virus injection before experiments were performed.

### Array Tomography

Mice injected with AAV(8)-CMV-DIO-Synaptophysin-YFP in EP were deeply anesthetized, perfused transcardially with room temperature phosphate-buffered saline (PBS) followed by 4% paraformaldehyde (PFA) in PBS. The brain was removed from the skull, post-fixed overnight at 4°C in 4% PFA, rinsed and stored in PBS. 250 µm thick coronal sections were cut with a Leica VT1000s vibratome. Sections containing the habenula with high Synaptophysin-YFP expression were noted using an epifluorescence microscope, and approximately 0.5 × 0.5 mm squares of tissue were cut out under a dissecting scope with Microfeather disposable ophthalmic scalpels. These small tissue squares were then dehydrated with serial alcohol dilutions and infiltrated with LR White acrylic resin (Sigma Aldrich L9774), and placed in a gel-cap filled with LR White to polymerize overnight at 50°C. Blocks of tissue were sliced on an ultramicrotome (Leica EM UC7) into ribbons of 70 nm sections.

Antibody staining of these sections was performed as described previously (Saunders et al. 2015). Briefly, antibodies were applied across multiple staining sessions (up to three antibodies per session) and a fourth channel left for DAPI. Typically, Session 1 stained against YFP (chicken α-GFP, GTX13970, GeneTex), Gephyrin (mouse α-Gephyrin, 612632, Biosciences Pharmingen), and Synapsin-1 (rabbit α-Synapsin-1, 5297S, Cell Signaling Tech); Session 2 for PSD-95 (rabbit α- PSD95, 3450 Cell Signaling Tech.); Session 3 for Vgat (mouse α-VGAT, 131 011 Synaptic Systems), and VGLUT2 (rabbit α-VGLUT2, 135 403 Synaptic Systems). In one sample the staining order was reversed, and revealed that order-dependent differences in staining quality did not alter the analysis. Each round of staining was imaged on a Zeiss Axio Imager upright fluorescence microscope before the tissue ribbons were stripped of antibody and re-stained for a new imaging session. Four images were acquired with a 63x oil objective (Zeiss) and stitched into a single final image (Mosaix, Axiovision). Image stacks were processed by aligning in Fiji with the MultiStackReg plug-in, first on the DAPI nuclear stain and with fine alignments performed using the Synapsin 1 stack. Fluorescence intensity was normalized across all channels, such that the top and bottom 0.1% of fluorescence intensities were set to 0 and maximum intensity, respectively.

Image analysis was performed as described previously (Granger et al. 2020). Pre-processing steps included trimming the image edges and masking out regions that correspond to cell nuclei as defined by DAPI signal. Background subtraction was performed at rolling ball radius of 10 pixels in Fiji with the Rolling Ball Background Subtraction plug-in. Synaptophysin-YFP channel was used to create 3D binary masks corresponding to EP *Sst+* terminals.

For co-localization analysis, antibody fluorescence puncta were fit with a gaussian distribution to identify and assign a pixel location corresponding to the centroid of the gaussian. The YFP mask was overlaid to the antibody puncta location distributions and co-localization was calculated as the number of pixels that overlapped within the YFP mask divided by the total number of pixels of the YFP mask. To estimate colocalization level by chance, the locations of each centroid were randomized prior to co-localization calculation. This randomization was repeated 1000 times to used calculate a Z-score per sample per antibody signal to pool across samples (Supplemental Figure 1).

For cross-correlation analysis, each antibody stack was z-scored and two stacks from the same sample were compared by shifted one image up to ± 10 pixels in increments of 1 pixel vertically and horizontally. At each shift, the co-variance of the images were calculated (Figure 1F). Co-variance was also measured specifically within the YFP mask by restricting the above calculation to the image area within the YFP mask (0.1∼0.6% of the total image) (Figure 1G).

### DMOS optical setup

A digital micromirror device (DMD) surface was exposed from a DLP LightCrafter Evaluation Module (Texas Instruments) and mounted in the optical path to direct the reflected laser beam to the back aperture of a 0.8 NA 40x objective lens (Olympus). A 473nm collimated beam of width ∼1mm was emitted from the laser (gem 473, Laser Quantum) and was uncollimated by passing through a static holographic diffuser (Edmund Optics) with 10° divergence angle. A mechanical shutter (Uniblitz, model LS6Z2) was mounted between the laser and the diffuser to control the timing of light exposure. The uncollimated, divergent light after the diffuser was converged using a lens (f = 30 mm) to cover the DMD surface. The diffracted beam from the DMD was collected by a second lens (f = 100 mm) and relayed to the back-aperture of the objective to form a conjugate DMD image in the sample plane. The optical setup achieved 22x magnification of the DMD image onto the sample plane with a resultant field of view of 299µm (width) x 168 µm (height).

Custom software written for ScanImage in MATLAB was used to control the individual DMD mirrors. Light power was controlled using Laser Quantum RemoteApp software via the RS232 port. The power efficiency of the system was ∼5% from laser output to specimen, resulting in maximum power of 10 mW at the sample plane when all mirrors were in the “on” position. The validation of the DMD alignment using electrophysiological recording was performed as shown in Supplemental Figure 3.

### Acute brain slice preparation

Acute brain slices were obtained from adult mice anesthetized by isoflurane inhalation and perfused transcardially with ice-cold, carbogen-saturated artificial cerebral spinal fluid (aCSF) containing (in mM): 125 NaCl, 2.5 KCl, 25 NaHCO_3_, 1.25 NaH_2_PO_4_, 2 CaCl_2_, 1 MgCl_2_, and 17 glucose (300 mOsm/kg). The brain was dissected, blocked, and transferred into a tissue slicing chamber containing ice-cold aCSF. 250-300 µm thick coronal slices containing LHb were cut using a Leica VT1000s or VT1200 vibratome. Following cutting, each slice was recovered for 9-11 min individually in a pre-warmed (34°C) choline-based solution containing (in mM): 110 choline chloride, 11.6 ascorbic acid, 3.1 pyruvic acid, 2.5 KCl, 25 NaHCO_3_, 1.25 NaH_2_PO_4_, 0.5 CaCl_2_, 7 MgCl_2_, and 25 glucose, then for at least 20 min in a secondary recovery chamber filled with 34°C aCSF. After recovery, the slices in aCSF were cooled down to and maintained at room temperature until use. Choline and aCSF solutions were under constant carbogenation (95% O_2_/5% CO_2_).

### Electrophysiology

For whole-cell recordings, individual slices were transferred to a recording chamber mounted on an upright customized microscope with the DMOS system. LHb neurons were visualized using an infrared differential interference contrast method under a 40x water-immersion Olympus objective. Epifluorescence (LED light source from X-Cite 120Q, Excelitas) was used to confirm virus expression and to identify regions displaying high density of *SSt+ tdTom+* axons within the LHb. Recording pipettes (2-3MΩ) were pulled from borosilicate glass using P-97 flaming micropipette puller (Sutter). Pipettes were filled with cesium-based internal recordings solution consisting of (in mM): 135 CsMeSO_3_, 10 HEPES, 1 EGTA, 4 Mg-ATP, 0.3 Na-GTP, 8 Na_2_-Phosphocreatine, 3.3 QX-314 (Cl-salt), pH adjusted to 7.3 with CsOH, and diluted to 290-295 mOsm/kg. Whole-cell voltage clamp recording was performed in acute slices continuously perfused with carbogenated aCSF at room temperature at a flow rate of 3∼4ml/min. After forming an intracellular seal with a target LHb neuron, 473nm light stimulus was delivered using the full field-of-view of the DMOS setup to activate oChIEF expressing *Sst+* presynaptic axons to confirm a synaptic transmission onto the postsynaptic cell. In LHb neurons that elicited PSCs, we subsequently delivered stimulation pulses (2∼5ms pulse duration, 100ms interstimulus interval) consisting of 96 patterns of 23×28 µm boxes that tiled the entirety of the DMOS field-of-view to identify regions that gave rise to PSCs due to groups of axons. Voltage-clamp recordings were amplified and low-pass filtered at 3 kHz using a Multiclamp700 B (Axon Instruments, Molecular Devices) and digitized at 10 kHz using an acquisition board (National Instrument). Data was saved with a custom version of ScanImage written in MATLAB with the DMOS package that enabled mapping of the electrophysiological recording that contain PSC elicited by photo-stimulation to a spatial coordinate on the sample plane. Using this mapping table, we were able to reconstruct a spatial heatmap indicating the location coordinate of pre-synaptic axons that synapsed onto the postsynaptic neuron that we recorded from. All recordings were performed with R,S-3-(2-carboxypiperazin-4-yl) propyl-1-phosphonic acid (CPP, 10μM Tocris) in bath solution to block NMDAR-mediated excitatory postsynaptic current.

For the compound PSC recording experiment described in Figure 3, LHb neurons were voltage-clamped at a holding potential of −70 mV while the DMOS system delivered a light stimulation pattern consisting of a spatiotemporal sequence of 96 different spots for five consecutive sweeps. The cell was subsequently depolarized to a holding potential of 0mV and delivered the same stimulation pattern for another five consecutive sweeps.

For the minimal stimulation PSC recording experiment described in Figure 4, LHb neurons were voltage-clamped at an intermediate holding potential of −35 mV or −27mV while the DMOS setup delivered light stimulation pattern of 96 different spots in each trial. To ensure that we are only targeting pre-synaptic boutons, tetrodotoxin (TTX, 1μM Tocris) and 4-Aminopyridine (4-AP, 400μM Tocris) were present in the bath solution at room temperature throughout the experiment. Initial five trials collected using high laser intensity were used to determine the spatial map of input-output responses in the recorded cell. Next, custom software written in MATLAB was used to select a few hotspots out of the 96 candidate spots to enable rapid collection of hundreds of trials of data in these hotspots. In some occasions, these spots were then subdivided into smaller regions and the final hotspots widths ranged from 10∼30 µm, depending on our ability to evoke a PSC after reducing the stimulation spot size. After finalizing a stimulation pattern, we then manually adjusted the laser intensity using the Laser Quantum RemoteApp software until some of these spots elicited PSCs stochastically upon repetitive stimulation.

For the serotonin perturbation experiment with DMD ring illumination (Figure 7A-B), LHb neuron voltage-clamp recordings were performed at holding potentials of −64mV and 10mV, in presence of CB_1_ receptor antagonist AM251 (1μM, Tocris) at physiological bath temperature. 1μM Serotonin hydrochloride (Tocris) was applied to perfusion chamber to compare the effect of serotonin on glutamate/GABA co-release at a macroscopic level. For the serotonin perturbation experiment with pre-synaptic terminal stimulation (Figure 7C-L), same experimental condition as in Figure 4 was used with 0.25μM serotonin hydrochloride (Tocris) to reduce synaptic release probability.

### Model simulations

We developed a biophysical model simulating a probabilistic neurotransmitter release with small variance in the vesicle content. To simulate excitatory and inhibitory postsynaptic currents due to a single vesicle release, we used the *alpha function* of the form:

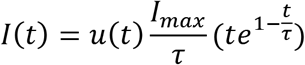

where *τ* is the time constant determining on- and off-kinetics of the function (τ_E_=1ms and τ_I_=3ms were used for excitatory and inhibitory PSCs, respectively), *I_max_* is the maximum amplitude of the current change, and *u*(*t*) is the impulse function that represents the onset of vesicle release. In the co-packaging version of the model, the excitatory and inhibitory PSCs occurred together and the vesicle noise was shared. In the independent version, the two PSCs occurred independently from each other with independent vesicle noise. The currents mediated by two different neurotransmitters were summed to generate net currents of two versions of release model:

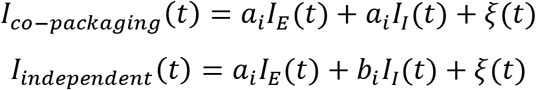

where *ξ*(*t*) is the white noise with standard deviation *σ* = 0.05, which scales with the signal size. *a*_*i*_ and *b*_*i*_ represent the scaling factor of the single vesicle content of the i^th^ trial

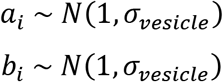

where *σ_vesicle_* is the standard deviation of fluctuations in the vesicle content across trials. We simulated hundreds of trials to generate a distribution of net currents using the same parameters for the two versions of model in MATLAB (available from https://github.com/seulah-kim/coreleaseAnalysis_Kim2021).

### Analysis of electrophysiology data

All analysis steps were performed in MATLAB (available from https://github.com/seulah-kim/coreleaseAnalysis_Kim2021). Schematic of analysis pipeline is shown in Supplemental Figure 4A.

#### Quality check

To ensure that we only include data collected with stable recording and that observed changes in evoked current peak size across trials are not due to variable amount of filtering due to fluctuations in resistance, access resistance between the pipette and the target cell was computed for every trial by fitting an RC response curve with two exponential functions and extrapolating the instantaneous peak size. The estimated access resistances across trials were median filtered, using window size of 2ms, to identify trials that exceeded 25% percentage of the initial access resistance, which was estimated from a median value of the first third trials of the total data recorded. In addition, we eliminated trials with >30pA drift in voltage-clamp recording within the trial. Across trials, any outliers that exceeded 30pA from the median of average trial value were eliminated.

#### Pre-processing

Raw current signals were baseline subtracted using the mean of baseline period (299.9 ms) of each trial. The offset signal was then low-pass filtered at 2kHz and smoothed using a savitsky-golay filter with polynomial order of 5 and frame length of 2.7 ms, followed by a moving median filter of 0.6 ms window. The current traces of all trials were grouped based on the stimulation location and then aligned with respect to the light onset of individual spots. Each trial was subsequently baseline offset based on the average current of the stimulation period.

#### Identification of putative hotspots and changepoint analysis

Median absolute deviation of individual time point was calculated across trials, for individual spots. If a spot contained time points that exceeded the 3 scaled median absolute deviation away from the median value for longer than five consecutive milliseconds, it was sorted as a hotspot. The rest of spots that did not meet these criteria were sorted as null spots. To determine the time window for trial-by-trial statistical analysis, change point analysis was performed on the light onset aligned traces of hotspots. This method identified an onset and an offset of evoked response time window such that the sum of the residual error of the three partitioned regions is minimized in the local root mean square level.

#### Fitting a noise model for individual cells

Null spots and 30ms period prior to photo-stimulation onset data were pooled to fit a gaussian distribution noise model for individual cells and extract standard deviation of the symmetric noise centered around the baseline current recording of each cell.

#### Maximum/minimum amplitude extraction and trial classification

In order to extract maximum and minimum amplitudes described in Figure 5, hotspots traces (time x trials) were further divided into pre-stim (−30ms to 0ms, relative to light-onset) and evoked periods. Maximum and minimum peak locations were identified trial-by-trial per hotspot for individual periods. Amplitudes of maximum and minimum peaks during evoked period were estimated by computing 1 ms average around the initial peak locations and subtracting the average value of the time window spanning -13ms to -3ms, prior to the individual peaks as baselines. Same steps were repeated using the pre-stim period data to create the null distribution of maximum and minimum amplitudes. Trials with either the maximum or minimum amplitude that was greater than 2 scaled standard deviation of symmetric noise of a given cell were classified as success. The rest of the trials were classified as failures.

#### Classification of hotspots and subtypes

To determine the final list of hotspots, we bootstrapped maximum and minimum amplitude pairs extracted from the pre-stim periods of individual hotspots 10,000 times to generate null distributions of probability of excitatory (p(E)), inhibitory (p(I)), and both (p∩ I) PSCs using the same criteria defined above for classifying trials as presence or absence of events. This was to account for spontaneous activity rate that would give rise to success rate observed during pre-stim period, and we wanted to ask whether observed success rate during the evoked period was statistically significant compared to the null success rate of pre-stim period.

Furthermore, we categorized individual hotspots into EPSC-only, IPSC-only, and both subtypes described in Figure 4. In EPSC only hotspots, only the p(E) during evoked period exceeded the 95% confidence interval of the bootstrapped null distribution of p(E). In IPSC only hotspots, only the p(I) of evoked period exceeded the 95% confidence interval of the bootstrapped null distribution of p(I). In both hotspots, both p(E) and p(I) of evoked period exceeded 95% CI of the bootstrapped null distributions of p(E) and p(I), respectively.

### Parametrization of model feature indicator

Model feature indicator derived from probability feature was computed by subtracting the probability value for which cdf=0.5 of p(E)*p(I) distribution (grey) from that of p(E ∩ I) distribution (purple) (Figure 5C,H and Figure 6A). For i_min_ feature output, model feature indicator was calculated as a difference in normalized minimum amplitude, i, for which cdf=0.5 between the groups with presence (solid red) and absence (dashed red) of an inhibitory current (Figure 5D,I). Similar analyses were performed for i_max_ feature output for maximum amplitudes between groups with presence (solid blue) and absence (dashed blue) of an excitatory current. Model feature indicators for correlation_all_ and correlation_s_ outputs were calculated as difference in correlation value for which cdf=0.5 between all trials (dark green) and shuffled (grey) and success only trials (light green) and shuffled (grey) groups, respectively (Figure 5E,J). For transformation of model feature indicator shown in Figure 6C, probability feature values less than 0 were assigned to zero and then normalized by 0.25, which is the theoretical maximum difference if p(E) and p(I) were assumed to be the same. Correlation features (correlation_all_ and correlation_s_) and cdf features (i_min_ and i_max_) values were cut off at 0 (floor) and 1 (ceiling). To reduce dimension after parametrization and transformation, we projected each spot on the model axis as the average of five model feature indicators (Figure 6D).

### Three types of noise metrics

Symmetric baseline recording noise was computed by fitting a gaussian function (mean and standard deviation) on pooled data consisting of portion of traces that are null spots (Supplemental Figure 4A) and 300ms baseline period across trials. Spontaneous activity peaks were extracted using the same method of minimum and maximum amplitude as described above applied to 30ms prior to photo-stimulation onset on each trial. Outlier fraction was calculated as the fraction of datapoints exceeding 3 scaled median absolute deviation from the pooled data consisting of null spots and 300ms baseline period.

### Analysis of 5-HT pharmacological effect

K-S test was performed with bootstrapping (10,000 times) with resampling size matching the smaller number of trials of the two groups (normally this is post 5-HT group size) to compare before and after 5-HT on the minimum and maximum amplitudes.

i_min_ and i_max_ subset distributions analysis (Figure 7M-N; Supplemental Figure 7D1-6) was performed by aligning individual trials by the i_min_ timepoint within the time window determined by changepoint analysis. Trials were sorted in ascending order based on the i_min_ size and then grouped in 10 trials. Maximum and minimum amplitudes were extracted from the average trace of each group aligned by i_min_ peak location.

For the prediction of model feature indicator change (Figure 7Q), the trials of pre 5-HT condition was analyzed with gaussian noise added to match the post 5-HT condition, subset of success trials were included to match the release probability of post 5-HT condition, and the i_min_ and i_max_ amplitudes of “both” success trials were scaled to match the scaling of pre vs. post 5-HT condition of median amplitudes of success trials.

### Statistical tests

Comparisons of proportions of hotspots were done using Fisher’s exact test. Bootstrapping (10,000 times) method was used to simulate variance in the sampling for statistical tests. Lower boundary of p-value for bootstrapped results was set by the bootstrap number (e.g. p = 1/10,000= 0.0001). Cumulative distributions were compared using Kolmogorov-Smirnov tests. P-values smaller than 0.001 were reported as p < 0.001.

## Acknowledgments

The authors thank Aurélien Bègue, Pavel Gorelik, and Ofer Mazor for help with the DMOS setup; Mahmoud El-Rifai and others for production and analysis of array tomography data; Adam Granger for assistance with array tomography data analysis; and Sabatini lab members for helpful discussions and feedback on the manuscript. This work was supported by NIH (R01NS103226 to B.L.S and P30NS072030 to the Neurobiology Imaging Facility). The authors declare no competing financial interests.

## Author Contributions

S.K., M.L.W., and B.L.S. designed electrophysiology experiments and discussed results. S.K. and B.L.S. built the DMOS system and developed computational models and analysis methods. S.K collected electrophysiology data, performed model simulations, intracranial injections, and histology, and analyzed all electrophysiology and array tomography data. M.L.W. and M.E.R. collected array tomography data. A.R.K. performed intracranial injections and histology. S.K. and B.L.S. wrote the manuscript with comments and feedback from the other authors.

## Tables

**Supplemental Table 1.**
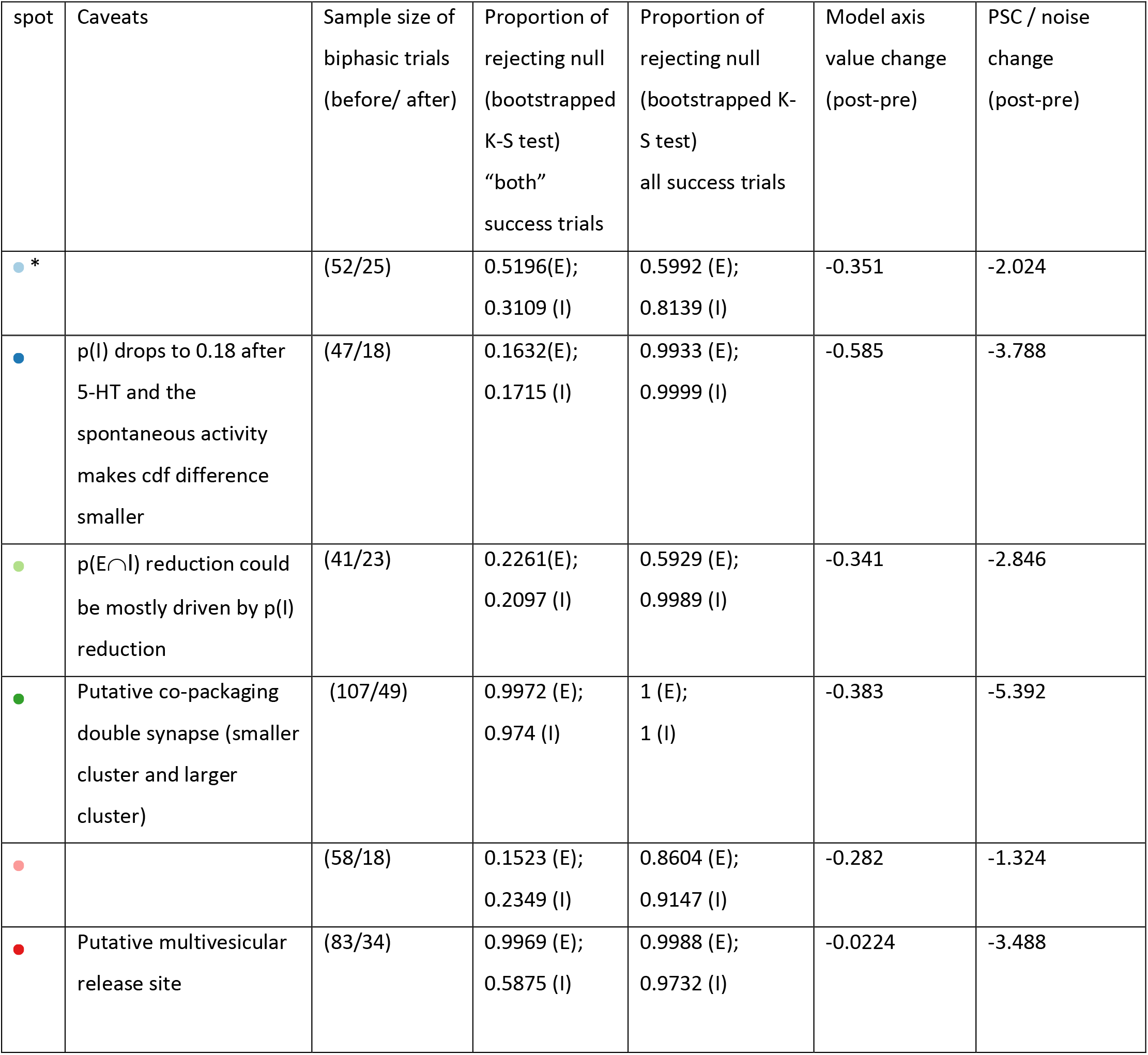
Summary of 5-HT effect on six example co-packaging uPSC sites. Related to Figure 7. Spot annotated with * corresponds to Figure7D-M

## Supplemental Figure Legends

**Supplemental Figure 1.**
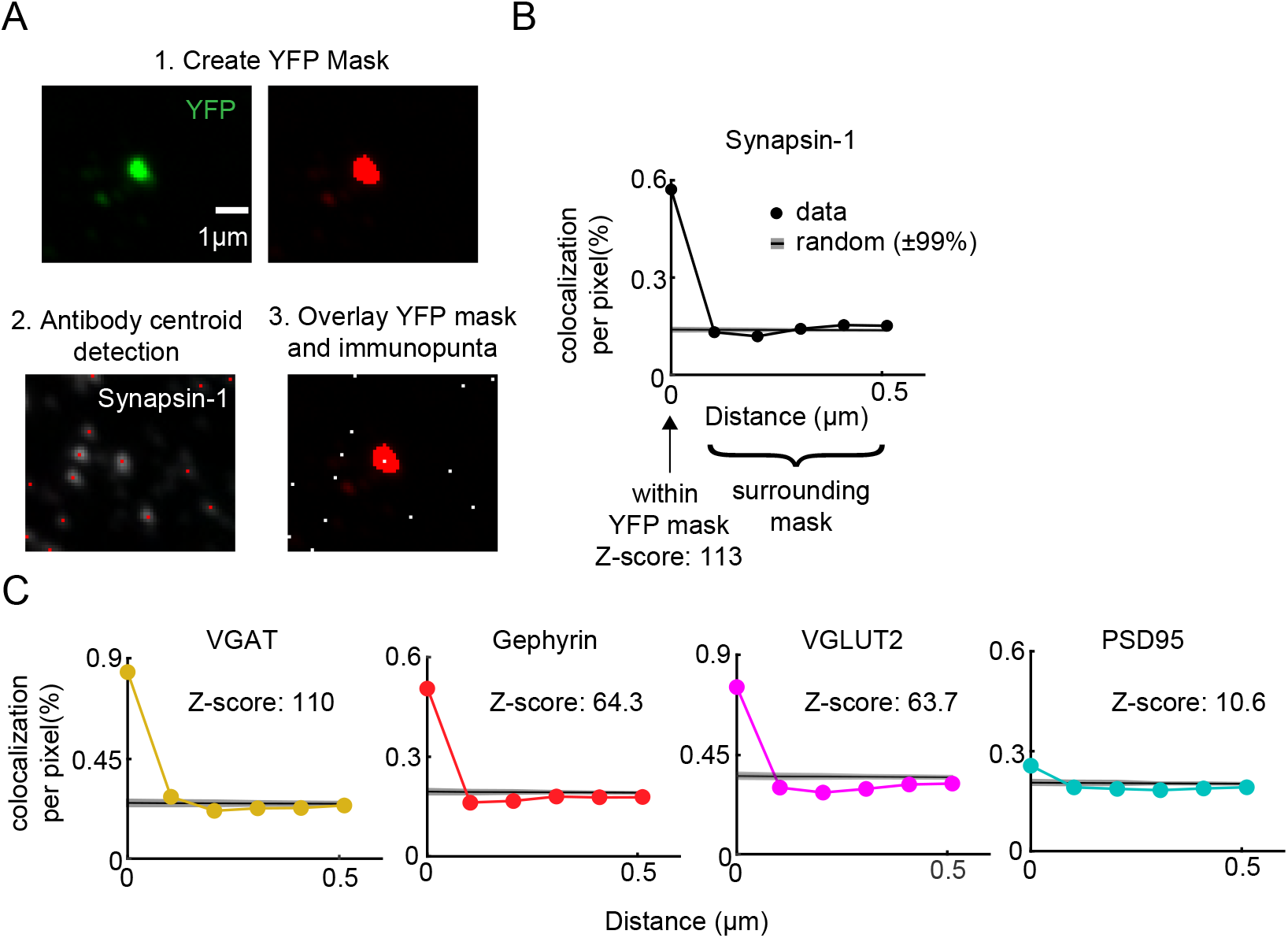
Co-localization analysis of antibodies within and outside of YFP-labelled EP Sst+ terminals. A) Co-localization analysis schematic. The YFP channel fluorescence was used to create masks to identify pixel regions containing labeled *Sst+* terminals. Each antibody channels was analyzed independently to extract the locations of the centroids of immunolabeled puncta. Extracted centroid locations were compared to the YFP masks in the same sample plane. For each immunolabeling channel, the percentage of pixels in the YFP+ masks that contained a punctum centroid was calculated and is referred to as the “co-localization” metric. B) Example synapsin-1 immunopuncta co-localization within the YFP mask and the surrounding regions compared to that expected by chance. Antibody locations were randomized 1000 times and the 99th percentile upper and lower boundaries are shown. Z-score is calculated as the difference between the mean antibody co-localization within the YFP mask and the mean randomized co-localization, divided by the standard deviation of the random co-localization. C) Example co-localization analysis results for Vgat, Gephyrin, Vglut2, and PSD95 antibodies from the tissue sample shown in B.

**Supplemental Figure 2.**
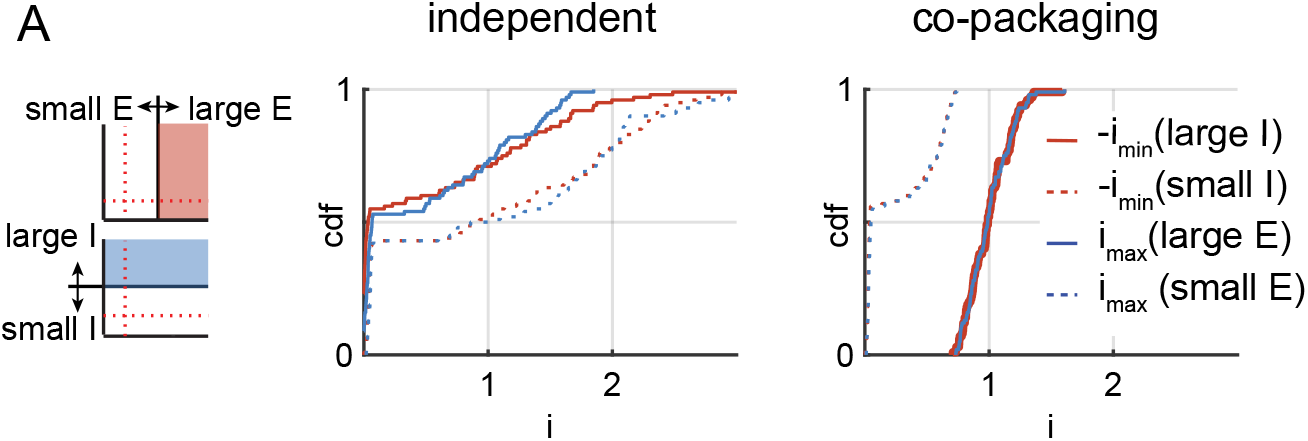
Accurate separation of synaptic failures is not required for the cdf analysis. A) Simulated cdfs of maximum PSC amplitudes (i_max_, blue) given a large (i_max_(E), solid) or small (i_max_(no E), dashed) excitatory current in the same trial for the independent (*left*) and co-packaging (*right*) release models. Similar analyses were performed for the minimum PSC amplitudes (-i_min_, red) given large (-i_min_(I), solid) or small (-i_min_(no I), dashed) inhibitory currents in the same trial. Simulation parameters are the same as in Fig 2D. Here we used the median to separate currents into large and small.

**Supplemental Figure 3.**
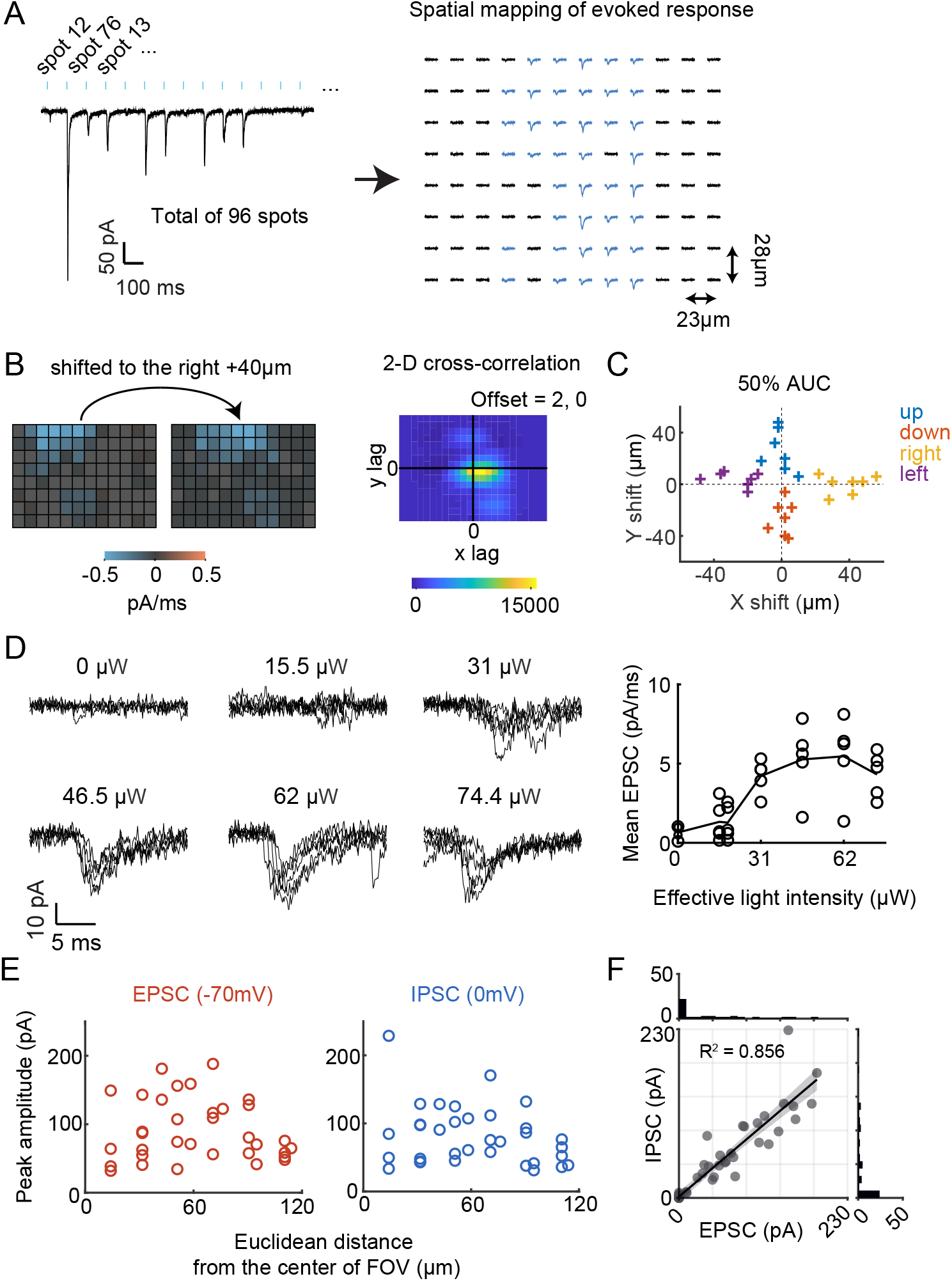
DMD based photo-stimulation enables spatially-specific activation of EP *Sst+* OChIEF-expressing terminals. A) *left*, Each trial consisted of rapid serial illumination of 23 x 28 µm photo stimulation spots to 96 different locations tiling the field of view in a pseudorandom spatial pattern such that the PSCs evoked from each spot are recorded in ∼10s voltage-clamp trace. Shown here is an example recording at −70 mV. *right,* The 96 PSCs in each trace are extracted and assigned to spatial locations based on the coordinates of the illuminated spots and the stimulus timing. B) Example showing that the DMD-evoked spatial map is consistent with the physical locations of EP *Sst+* terminals. *left,* the input-output relationship was initially mapped (V_h_=-70mV) using the DMD-based optogenetic stimulation platform then the brain slice was shifted in +x direction relative to the microscope objective lens by 40 µm after which the input-output relationship was re-mapped. *right*, 2D cross-correlation of the two spatial maps (before and after the objective lens movement) reveals that the two images are offset by 2 pixels, as predicted by the pixel spacing. The offset is calculated from the X and Y locations where the cumulative sums of correlation coefficient across y and x, respectively, reach the 50% of the total sum. Spatial maps are calculated from the average of 5 trials. C) Summary of quantification of cross-correlation calculated shifts as described in (B) for 7 cells from 3 animals. Colors indicate the direction of the slice movement. D) Saturation of the amplitude of the evoked PSC from the same 23 x 28 µm photo-stimulation spot in an example LHb neuron. *left,* electrophysiological recording (V_h_=-70 mV) for 5 trials at each indicated light intensity. Traces correspond to 5-25 ms time window after stimulation onset. *right,* Individual (circle) and average (line) EPSC amplitudes as a function of illumination intensity. E) Relationship between distance of the stimulation spots from the LHb cell body (located at the center of field of view (FOV)) and the corresponding evoked EPSC (*left*) and IPSC (*right*) peak amplitudes (data shown for the same neuron as in Fig 3F). F) Scatterplot of IPSC vs. EPSC peak amplitude pairs evoked at photo-stimulated spots within 80 µm perimeter from the center of the field of view from the neuron analyzed in Fig 3F.

**Supplemental Figure 4.**
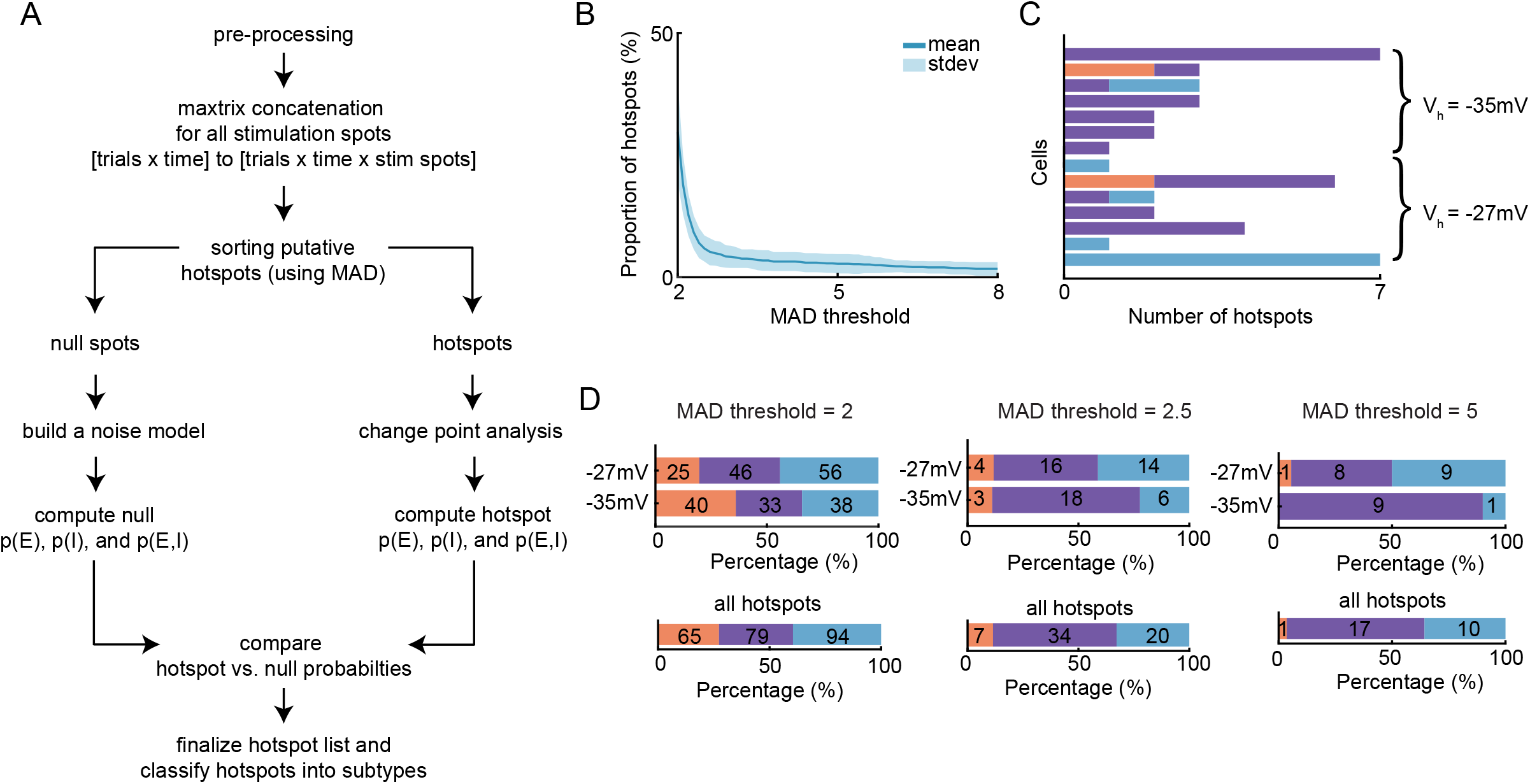
Automated analysis of evoked unitary responses. A) Hotspot detection and classification analysis pipeline flowchart (see Methods). B) Effect of median absolute deviation (MAD) threshold on the proportion of putative hotspots out of total stimulation spots. The MAD threshold, expressed in multiples of the empirically measured MAD for each cell, determines the selection of putative active hotspots which are required to have current deviation that exceed the threshold at least 5ms (the branching step in panel A). Mean and standard deviation of the proportion of illuminated spots designated as hotspots (data from 14 cells are shown). A MAD threshold of 3 was used for Figure 4F. C) Distribution of putative hotspot numbers across all cells (n=14 cells, 9 animals). MAD threshold of 3 was used. The holding potential of individual cells is indicated. PSCs are designated as EPSCs only (red), IPSCs only (blue), or both (purple). D) Effect of MAD threshold on the proportion of final hotspot subtypes. As in Figure 4F for MAD threshold of 2 (*left*), 2.5 (*middle*), and 5 (*right*). Color code as in panel (C).

**Supplemental Figure 5.**
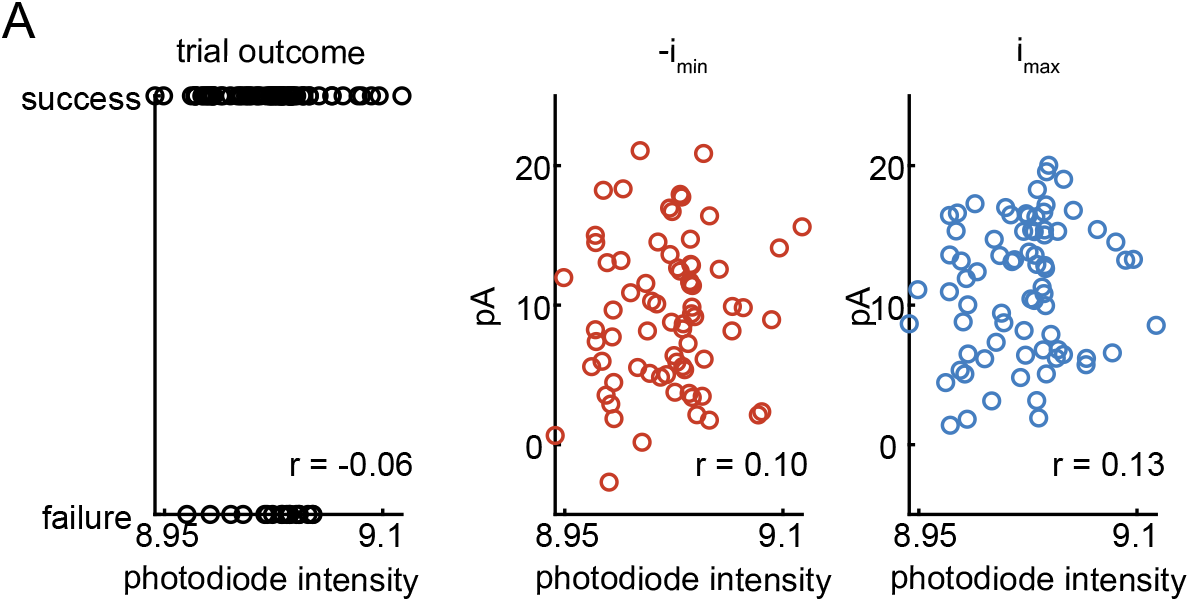
Unitary response correlation is not driven by stimulus fluctuations. A) Stimulation intensity fluctuation as detected by a photodiode versus trial-by-trial outcome (left) and amplitudes of -i_min_ and i_max_. Same dataset as in Figure 5F-J.

**Supplemental Figure 6.**
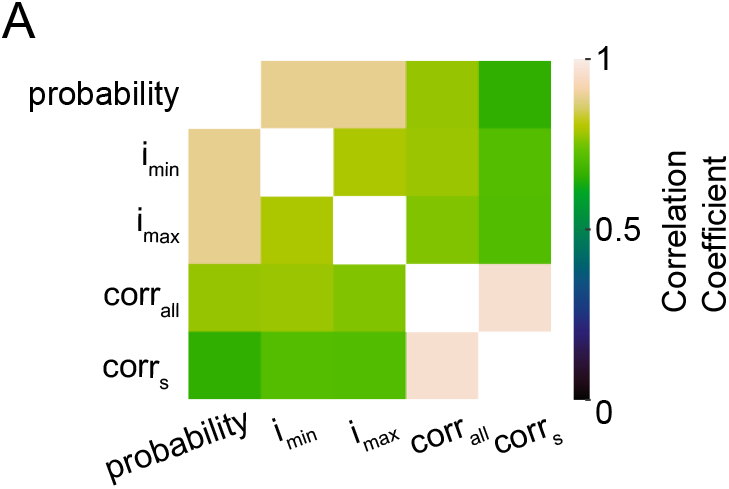
Model feature indicators are correlated in their support for the co-packaging model. A) Correlation heatmap of model feature indicators. Color represents the pair-wise correlation of each model feature across the same spots in (C).

**Supplemental Figure 7.**
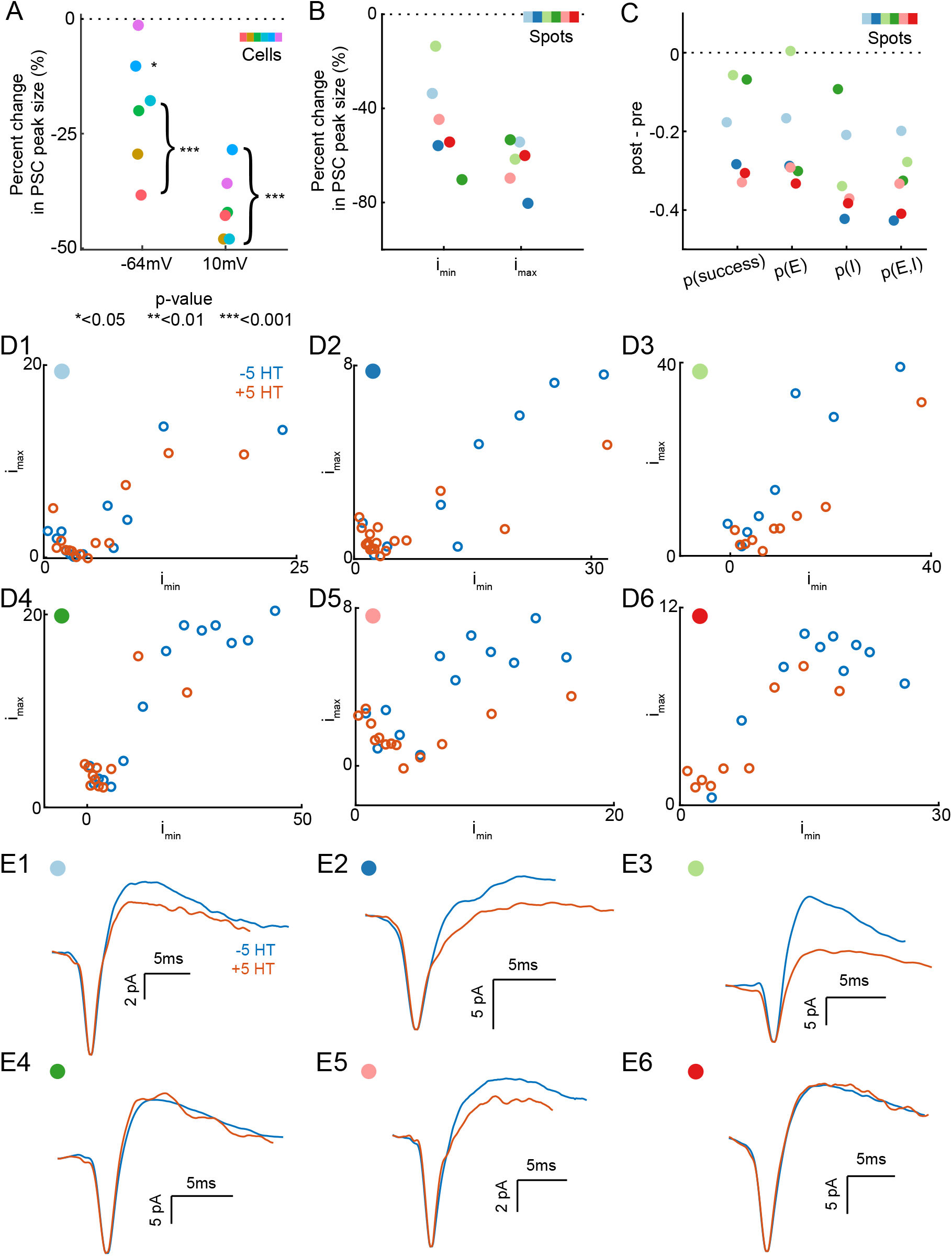
5-HT reduces probabilities of detecting EPSCs and IPSCs evoked by EP *Sst*+ terminal activation. A) Peak amplitude changes in the DMD ring stimulation evoked in the composite EPSC (−64 mV) and IPSC (10 mV) as result of 5-HT bath application. Each dot represents the difference in mean evoked peak amplitude of 15 trials before and after 5-HT application. Asterisks represent significance level of unpaired t-test comparing pre and post 5-HT groups. Colors indicate cell identity. B) Average relative minimum and maximum amplitude changes of DMOS-evoked unitary biphasic spots across all trials as result of 5-HT bath application. Colors indicate spot identity consistent as in Figure 7O. C) Changes in probabilities of detecting success trials, EPSC, IPSC, and both trials due to 5-HT bath application for DMOS-stimulated unitary biphasic spots. Each dot represents the difference in probabilities calculated from scatterplot of each spot before and after 5-HT. Colors and markers are consistent as in panel B. D) The effect of 5-HT on subset distributions of minimum and maximum amplitudes of co-packaging sites, without sorting trials by success and failures. Scatter corresponds to the amplitudes of the average trace of different subsets of dataset before (blue) and after (red) 5-HT bath application. Each dot in the top right indicates spot identity consistent as in Figure 7O. E) The effect of 5-HT on the average waveform of co-packaging sites, without sorting trials by success and failures. Average of each trial was aligned by the minimum peak location within the analysis time window. Before (blue) and after (red, normalized by the minimum peak amplitude of “before” condition) 5-HT bath application traces are compared. Each dot in the top right indicates spot identity as in Figure 7O.

